# Unique mineralization pattern revealed in TBCK syndrome mouse model

**DOI:** 10.64898/2026.02.18.706703

**Authors:** Kaitlin A. Katsura, Yuchen Jiang, Marius Didziokas, Nir Z. Badt, Sonia Dougherty, Kyle H. Vining, Elizabeth J. Bhoj

## Abstract

TBCK syndrome is a severe degenerative leukoencephalopathy with multisystem involvement. Neurodevelopmental, craniofacial, and pulmonary challenges are among the topmost effects on these children. TBCK has been implicated in endo-lysosomal regulation, RNA transport, and mTOR-associated pathways, all of which are critical for the development of mineralized tissue. Although craniofacial abnormalities can be clinically apparent, conventional imaging approaches may overlook subtle defects in mineral quality.

Here, we apply our multimodal framework to investigate the mineralization of enamel, dentin, and alveolar bone in a *Tbck* knockout mouse model. This is the first time our multimodal framework will be applied to a genetic condition. Using micro-computed tomography (microCT), histology, nanoindentation, energy-dispersive spectroscopy, and Raman spectroscopy, we identify tissue- and stage-dependent mineral effects undetected by microCT alone. *Tbck* loss resulted in differences in enamel and dentin element compositions as early as secretory and transition stages, while mechanical properties remained undetected until maturation stage.

Notably, *Tbck* knockout enamel exhibited reduced calcium and phosphorus content, along with increased carbon content during early mineralization, consistent with the retained organic matrix. Additionally, marked and opposing alterations in magnesium and iron levels began at the secretory stage. Together, these findings define a previously unrecognized mineralization signature associated with TBCK deficiency and establish multimodal hard-tissue analysis as a sensitive approach for detecting early craniofacial phenotypes in rare genetic disorders.

## Introduction

### TBCK syndrome is a rare and severe neurodevelopmental disorder characterized by profound hypotonia, developmental delay, seizures, and progressive multisystem involvement^1–8^

Although skeletal abnormalities and craniofacial differences are clinically apparent in the majority of affected children, the timing, tissue specificity, and underlying mineralization defects associated with this disease remain poorly defined. In particular, the oral and dental manifestations of TBCK syndrome have not yet been systematically described.

Children with TBCK syndrome are reported to exhibit early-onset osteoporosis^4,8,9^, suggesting impaired mineral quality rather than gross defects in skeletal patterning. Craniofacial differences are also one of the most frequently noted characteristics, yet whether these arise from altered bone formation, dental development, or both is unknown^10^. Often, these children have increased susceptibility to tooth aspiration and subsequent pulmonary complications. Despite the clinical relevance of dentition to feeding, airway protection, and infection risk, dental hard tissues have not been examined as a target organ system in TBCK syndrome.

Teeth provide a unique biological record of prenatal and early childhood development. Exfoliated and developing teeth have been widely proposed as non-invasive indicators of early-life environmental perturbations due to their features of incremental deposition in dentin and irreversible mineralization of both dentin and enamel. Among mineralized tissues, enamel is particularly distinctive, as it does not undergo remodeling and is formed through a tightly regulated, stage-specific process. During the secretory state, ameloblasts deposit the full thickness of enamel matrix proteins, followed by a brief transition phase and a prolonged maturation stage during which mineral content increases dramatically through matrix removal and hydroxyapatite crystal growth. These features make enamel especially sensitive to disruptions in cellular trafficking, matrix processing, and mineral transport.

At the molecular level, TBCK has been linked to RNA transport, endo-lysosomal function, and pathways linked to mTOR signaling, processes that are central to protein trafficking, matrix turnover, and mineral homeostasis^7,11–16^. Disruption of these pathways would be expected to disproportionately affect tissues such as enamel and bone, which rely on the precise coordination of secretion, degradation, and mineral deposition. However, whether TBCK deficiency alters craniofacial hard-tissue mineralization, and at what developmental stages such effects may arise, has not been investigated.

The continuously erupting mouse incisor provides a powerful model system for addressing these questions, as it contains spatially resolved zones corresponding to secretory, transition, and maturation stages of enamel and dentin formation within a single tissue. In recent work, we established a multimodal analytical framework combining micro–computed tomography, histology, mechanical testing, and spectroscopic approaches to characterize stage-specific mineral properties of craniofacial hard tissues. While this approach revealed striking developmental changes in enamel composition and mechanics, it has not yet been applied to a genetic disease model.

Here, we apply this multimodal framework to a *Tbck* knockout mouse model to test the hypothesis that TBCK deficiency produces subtle but biologically meaningful alterations in craniofacial hard-tissue mineralization that are not captured by conventional imaging alone^17^. We demonstrate that standard techniques, such as microCT and histology, may underestimate disease-associated phenotypes, whereas integrated mechanical and compositional analyses reveal early and tissue-specific mineral defects during tooth development. Using TBCK syndrome as a model, this study highlights the sensitivity of enamel and bone as readouts of genetic perturbations affecting early craniofacial development.

## Results

### TBCK deficiency alters incisor mineralization dynamics detectable by microCT

To assess whether global *Tbck* loss affects craniofacial hard-tissue mineralization, we first performed micro–computed tomography (microCT) on mandibles from wild-type (WT), *Tbck* heterozygous (Het), and *Tbck* knockout (KO) mice. Qualitative inspection revealed subtly altered mineralization patterns along the continuously erupting incisors and molars in *Tbck* KO mice compared with WT controls (Fig. 1a).

**Fig. 1.**
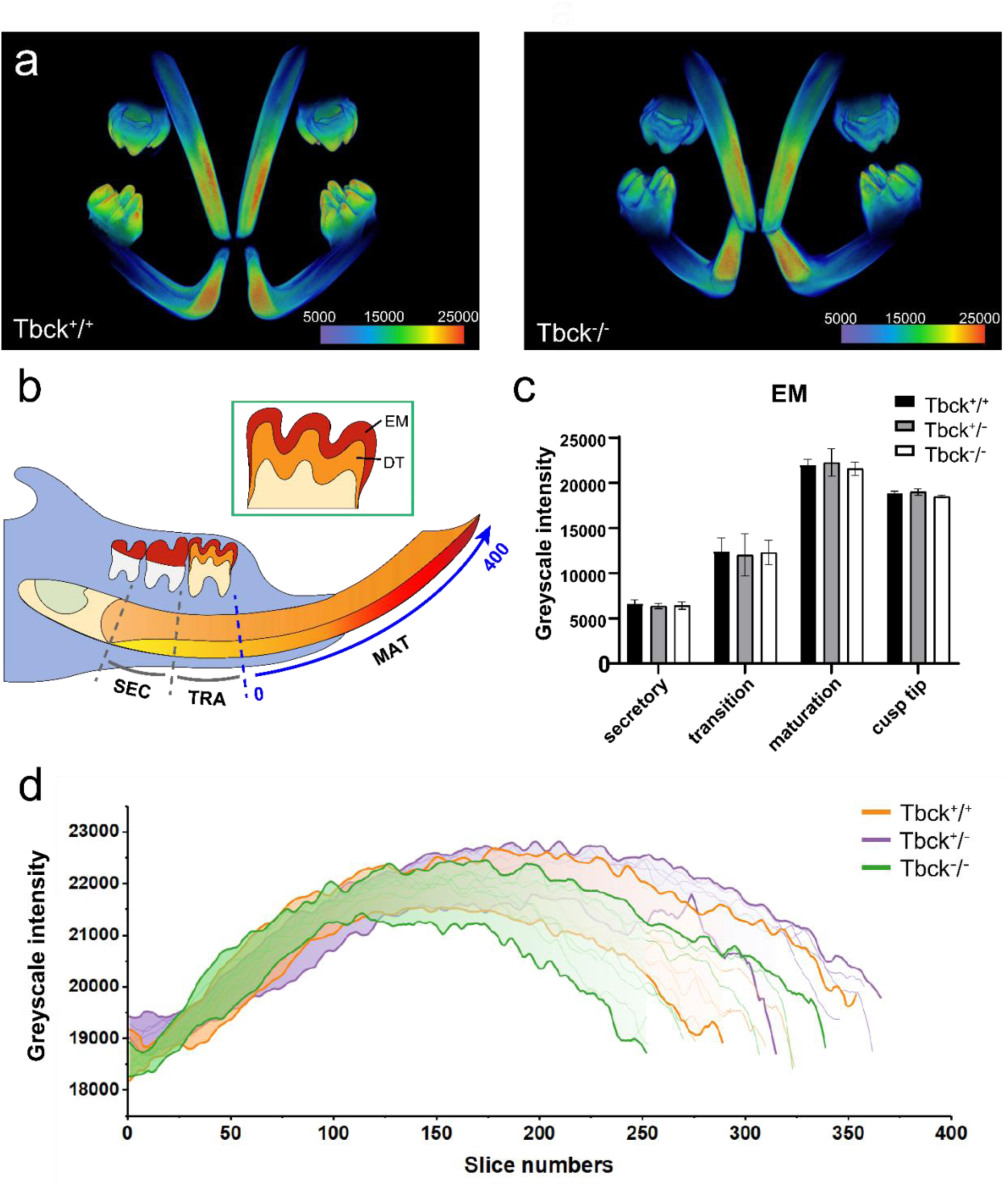
TBCK deficiency alters incisor mineralization dynamics assessed by micro–computed tomography (microCT). **a** Representative microCT reconstructions of mandibles from Tbck^+/+^ (WT) and Tbck^−/−^ (KO) mice, displayed as pseudocolor grayscale intensity maps. **b** Schematic of the continuously erupting incisor illustrating the secretory (SEC), transition (TRA), and maturation (MAT) regions and the sagittal section strategy used to generate grayscale intensity profiles in **d**. The inset indicates enamel (EM) and dentin (DT) compartments used for data collection in the molar. **c** Quantification of enamel grayscale intensity in cropped regions of interest defined for incisor enamel and molar enamel^17^. Data are presented as mean ± SD; statistical comparisons were performed using 2-way ANOVA with significance defined as p < 0.05. **d** Enamel sagittal grayscale intensity profiles along the incisor for Tbck^+/+^, Tbck^+/−^, and Tbck^−/−^ mice. Measurements started at slice 0 were defined as the position at which enamel and dentin exhibited equal grayscale intensity (see panel **b**). n=6 per genotype.

To enable standardized comparison across samples, sagittal sections were aligned using a grayscale-based normalization strategy, defining a reference slice corresponding to equivalent enamel and dentin intensity (Fig. 1b). While overall enamel grayscale intensity did not differ significantly between genotypes, *Tbck* KO incisors exhibited a trend of reduced mineralized length and decreased total incisor volume (Fig. 1c–d, S1). Divergence in grayscale profiles became apparent during the eruption phase, coinciding with the transition from alveolar bone encasement to functional exposure.

In contrast, molar mineral density and volume were largely preserved across genotypes, although heterozygous females displayed increased molar root length relative to WT and KO controls (Fig. S1). Together, these findings suggest that TBCK deficiency alters the timing and spatial progression of incisor mineralization rather than gross mineral density.

### Enamel matrix organization is disrupted during the transition and maturation stages in *Tbck* knockout mice

Given the temporal divergence observed by microCT, we next examined enamel matrix organization using sagittal histological sections stained with hematoxylin and eosin (H&E). In WT incisors, the enamel matrix exhibited the expected orderly progression from secretory to transition to maturation stages.

In *Tbck* KO mice, enamel matrix disruption was first evident during the transition stage and became more pronounced in the maturation zone (Fig. 2a, b2, c2). The affected regions exhibited irregular enamel matrix appearance most proximal to the dentin enamel junction, when compared with WT controls. These histological changes present at an earlier time point than what was detected by microCT, exhibiting altered mineralization dynamics at the extracellular protein level.

**Fig. 2.**
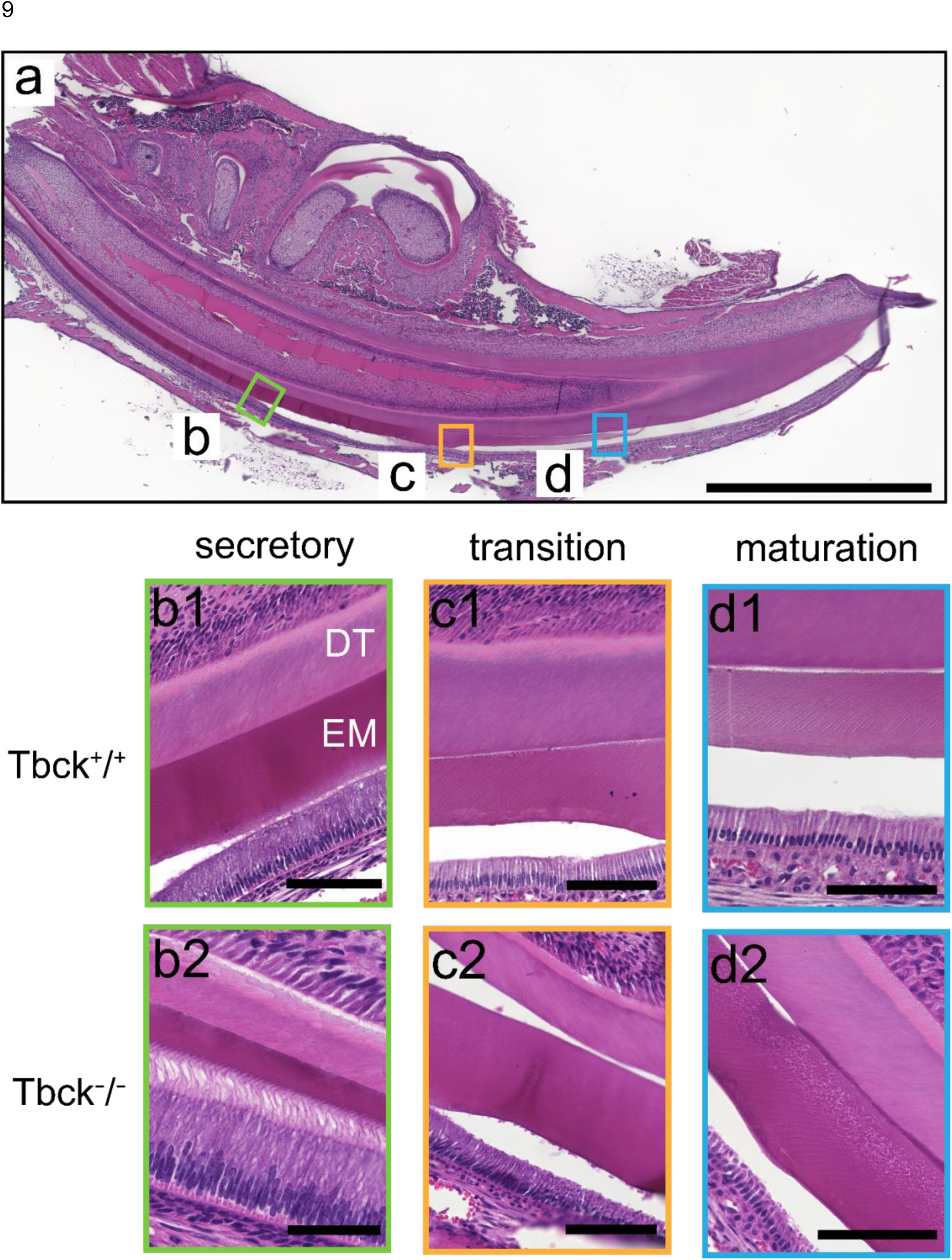
H&E histology reveals disrupted enamel matrix organization in Tbck knockout incisors during transition and maturation. Representative sagittal sections of mouse mandibles stained with hematoxylin and eosin (H&E) illustrate incisor developmental regions. **a** Low-magnification overview of a Tbck^+/+^ section with boxes indicating locations imaged at higher magnification. Scale bar, 400 μm. **b–d** Higher-magnification views of secretory (b), transition (c), and maturation (d) regions showing enamel (EM) and dentin (DT) in Tbck^+/+^ (b1–d1) and Tbck^−/−^ (b2–d2) incisors. Scale bars, b1(80 μm), b2-d1(100 μm), d2(50 μm).

### TBCK deficiency reduces enamel mechanical properties at late mineralization stages

Motivated by dental anomalies noted by clinicians caring for individuals with TBCK syndrome, we next quantified the mechanical properties of craniofacial hard tissues by nanoindentation. Nanoindentation was performed on enamel (EM) and dentin (DT) across defined incisor enamel developmental regions (secretory, transition, and maturation) and in unerupted developing first molars (cervical crown and cusp tip), with additional measurements acquired in alveolar bone anterior to the first molar (Fig. 3a–d) ^17^. While enamel modulus and hardness were comparable between genotypes at earlier stages, *Tbck* KO enamel exhibited significantly reduced modulus and hardness at later stages of mineralization, with the most significant differences observed in incisor maturation stage and at cusp tip (Fig. 3a, c).

**Fig. 3.**
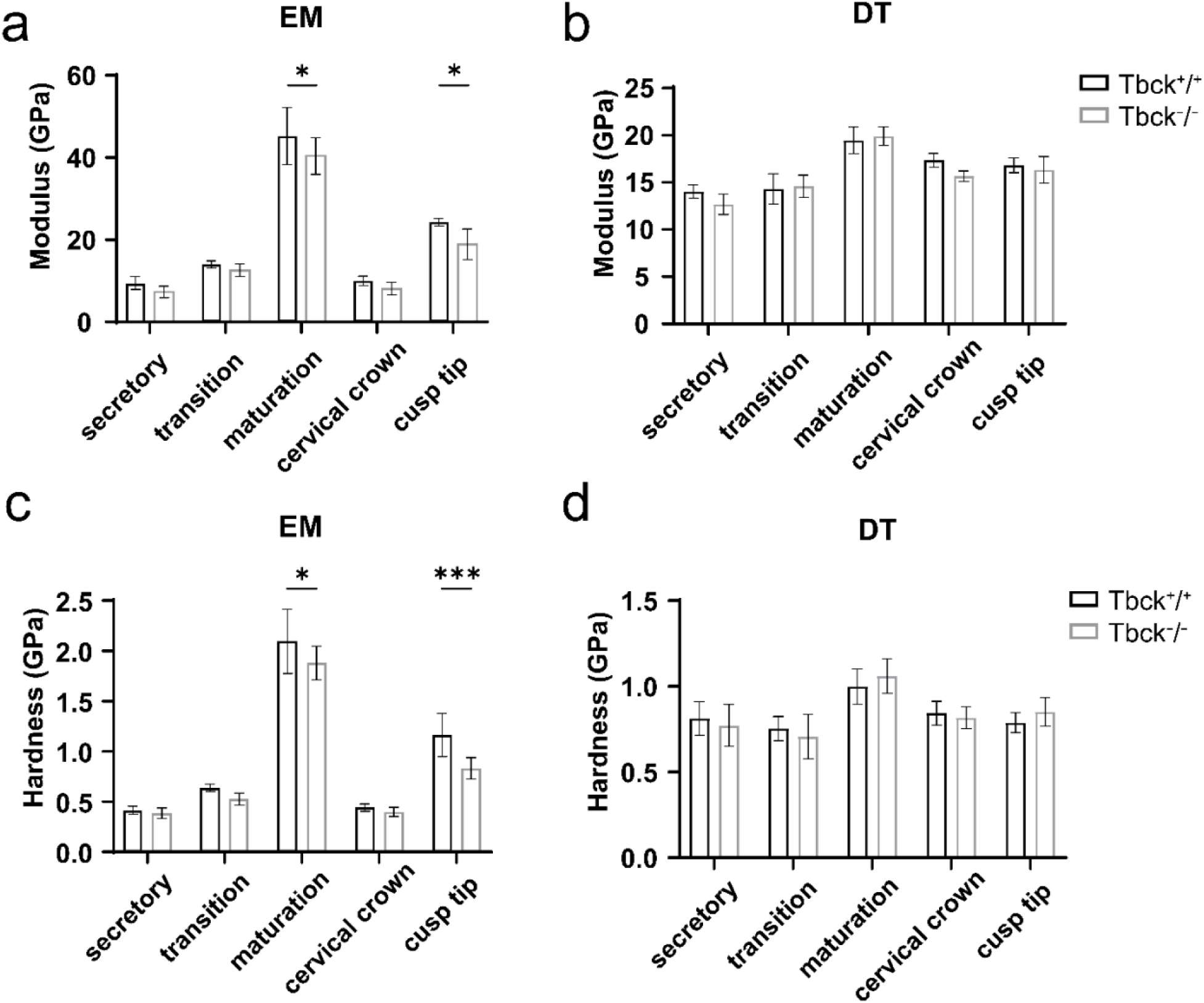
Nanoindentation identifies reduced enamel modulus and hardness in Tbck knockout mice at late mineralization stages. Enamel (EM) and dentin (DT) mechanical properties were quantified by nanoindentation in incisor (secretory, transition, maturation) and molar (cervical crown, cusp tip) for Tbck^+/+^ and Tbck^−/−^. **a–b** Elastic modulus (E) of enamel (a) and dentin (b). **c–d** Indentation hardness (H) of enamel (c) and dentin (d). Data are presented as mean ± SD; mice of both sexes were included (n = 6 indent sites per region). Statistical comparisons were performed using 2-way ANOVA; \**p* < 0.05, \*\*\**p*< 0.001.

In contrast, dentin mechanical properties were largely preserved, with no *Tbck*-dependent differences in dentin modulus or hardness across regions in either incisor or molar measurements (Fig. 3b, d). Consistent with skeletal findings reported in TBCK syndrome^10,18^, nanoindentation of alveolar bone also revealed significant reductions in both modulus and hardness in *Tbck* KO mice (Fig. S3). Together, these findings suggest that TBCK deficiency preferentially impacts late-stage enamel matrix hardening, while dentin mechanics remain largely intact.

### Elemental composition of enamel and dentin is altered early in mineralization in *Tbck* knockout mice

Energy dispersive X-ray spectroscopy (EDS) was performed to quantify atomic percentages of carbon (C), calcium (Ca), phosphorus (P), magnesium (Mg), and iron (Fe) across defined developmental regions of enamel (Fig. 4a–c) and dentin (Fig. S4a–b), with additional measurements in alveolar bone (Fig. S5).

**Fig. 4.**
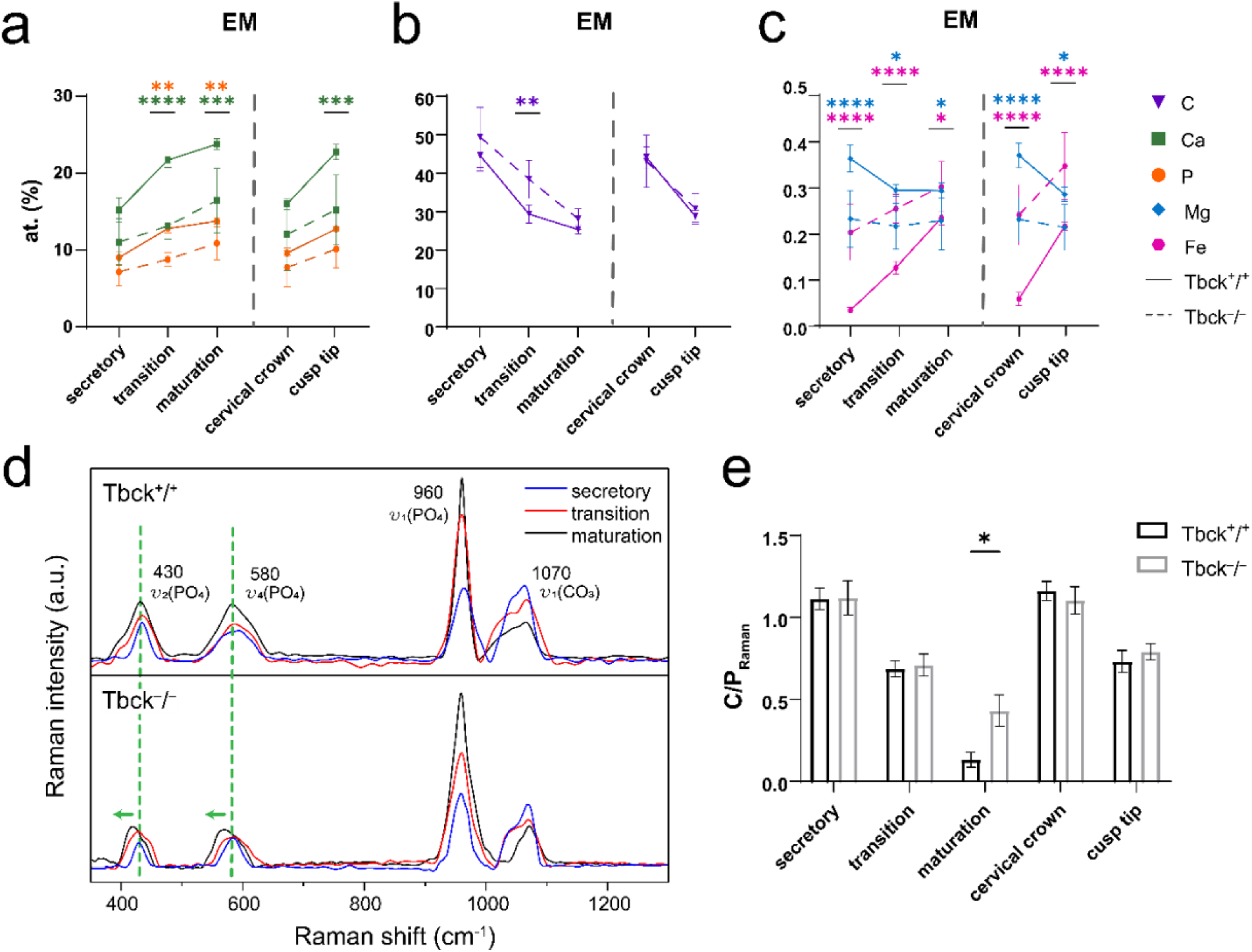
Tbck knockout (Tbck-/-) alters enamel elemental contents and late-stage chemical compositions assessed by EDS and Raman spectroscopy. **a** Atomic percent of calcium (Ca) and phosphorus (P) across regions. **b** Atomic percent of carbon (C). **c** Atomic percent of minor elements Mg and Fe. **d** Average Raman spectra from incisor enamel for Tbck^+/+^ (top) and Tbck^−/−^ (bottom) showing phosphate-associated bands ν_2_(PO_4_^3-^) (430 cm⁻¹), ν_4_(PO_4_^3-^) (580 cm⁻¹), and ν_1_(PO_4_^3-^) (960 cm⁻¹), as well as the carbonate ν_1_(CO_3_^2-^) band (1070 cm⁻¹); dashed markers indicate phosphate band positions used to compare genotypes. **e** Raman carbonate-to-phosphate ratio (C/P) of enamel is calculated as the integrated area of the carbonate band (∼1070 cm^−1^) divided by the phosphate band (∼960 cm^−1^). Data are presented as mean ± SD (n = 6 per genotype). \**p* < 0.05.

In enamel, Ca and P increased along incisor development and were higher at molar cusp tip than cervical crown (Fig. 4a). In both incisor and molar, *Tbck* KO enamel exhibited reduced Ca and P levels relative to Tbck WT, with significant differences evident at transition and maturation stages and molar cusp tip (Fig. 4a). Carbon decreased along enamel development (Fig. 4b), with the most considerable genotype difference observed at transition stage. Minor element profiles showed that Mg declined from secretory to maturation enamel and from cervical crown to cusp tip (Fig. 4c), and Mg level was consistently lower in *Tbck* KO enamel in incisor and molar. In contrast, Fe at% increased with enamel development in both genotypes, and *Tbck* KO enamel showed higher Fe at% beginning at secretory stage and persisting through transition and into molar regions, including a significant increase at cusp tip (Fig. 4c).

In dentin, C level was comparatively stable along incisor, with a modest increase in *Tbck* KO most apparent at maturation stage (Fig. S4a). Ca and P levels in dentin showed more minor developmental changes than enamel but were reduced in KO, with significant differences emerging from transition to maturation along incisor and at molar cusp tip (Fig. S4a). Mg at% in dentin also differed by genotype across development, with lower Mg levels in KO incisors and molars (Fig. S4b). Fe was elevated in *Tbck* KO dentin across incisor stages and in molars (Fig. S4b). In alveolar bone, *Tbck* KO mice exhibited reduced Ca and P levels compared with WT, while C level was comparable between genotypes (Fig. S5). Among minor elements, *Tbck* KO bone showed reduced Mg and increased Fe (Fig. S5).

### TBCK deficiency alters mineral signatures during late-stage mineralization

Given the compositional differences detected by EDS, we next evaluated mineral chemistry by Raman spectroscopy across matched developmental regions^10,19^. In incisor enamel, spectra from *Tbck* WT and KO samples exhibited the expected phosphate-associated vibration modes, including ν2(PO43-) (430 cm⁻¹), ν4(PO43-) (580 cm⁻¹), and ν1(PO43-) (960 cm⁻¹), as well as the carbonate ν1(CO32-) band (1070 cm⁻¹) (Fig. 4d)^20^. In Tbck WT enamel, the 960 cm⁻¹ phosphate peak increased from secretory to maturation. In contrast, *Tbck* KO enamel showed a more stepwise increase, with 960 cm⁻¹ peak intensity at transition stage remaining closer to secretory and a larger change occurring between transition and maturation (Fig. 4d). Across stages, the relative carbonate contribution decreased with maturation in both genotypes; however, *Tbck* KO enamel exhibited a higher carbonate-to-phosphate (C/P) ratio (CO32-/PO43-; integrated area of 1070 cm⁻¹ divided by 960 cm⁻¹) at maturation stage compared with WT (Fig. 4e). In addition, the ν2 and ν4 phosphate bands were shifted in *Tbck* KO enamel relative to WT, with the shift most evident in maturation spectra (Fig. 4d, green markers).

In molar enamel, spectra were similar between genotypes across cervical crown and cusp tip (Fig. S6). In both WT and KO samples, cusp tip exhibited a higher 960 cm⁻¹ phosphate peak than cervical crown, with a corresponding lower carbonate-to-phosphate ratio at cusp tip. No Tbck-dependent differences were detected in molar enamel C/P ratio (Fig. 4e).

In dentin, Raman spectra showed prominent phosphate and carbonate bands in incisor and molar (Fig. S7). Along the incisor development, both genotypes exhibited a progressive increase in 960 cm⁻¹ phosphate peak intensity from secretory to maturation with a concomitant decrease in carbonate contribution, reflected by declining C/P ratios across development (Fig. S7a; Fig. S8). Notably, *Tbck* KO dentin exhibited a significantly higher C/P ratio at incisor maturation stage and cusp tip compared with WT (Fig. S8).

### Integration of multiscale hard-tissue metrics using PCA and genotype prediction by MLR

To integrate microCT greyscale intensity with mechanical and compositional measurements, we applied a two-step multivariate workflow comprising principal component analysis (PCA) for dimensionality reduction and multiple linear regression (MLR) for genotype prediction^17^. Enamel and dentin were analyzed separately using the same set of variables: hardness, EDS/Mg (at%), EDS/Fe (at%), EDS/Ca (at%), EDS/P (at%), greyscale intensity, and the carbonate-to-phosphate ratio (C/P) (Fig. 5a–f).

**Fig. 5.**
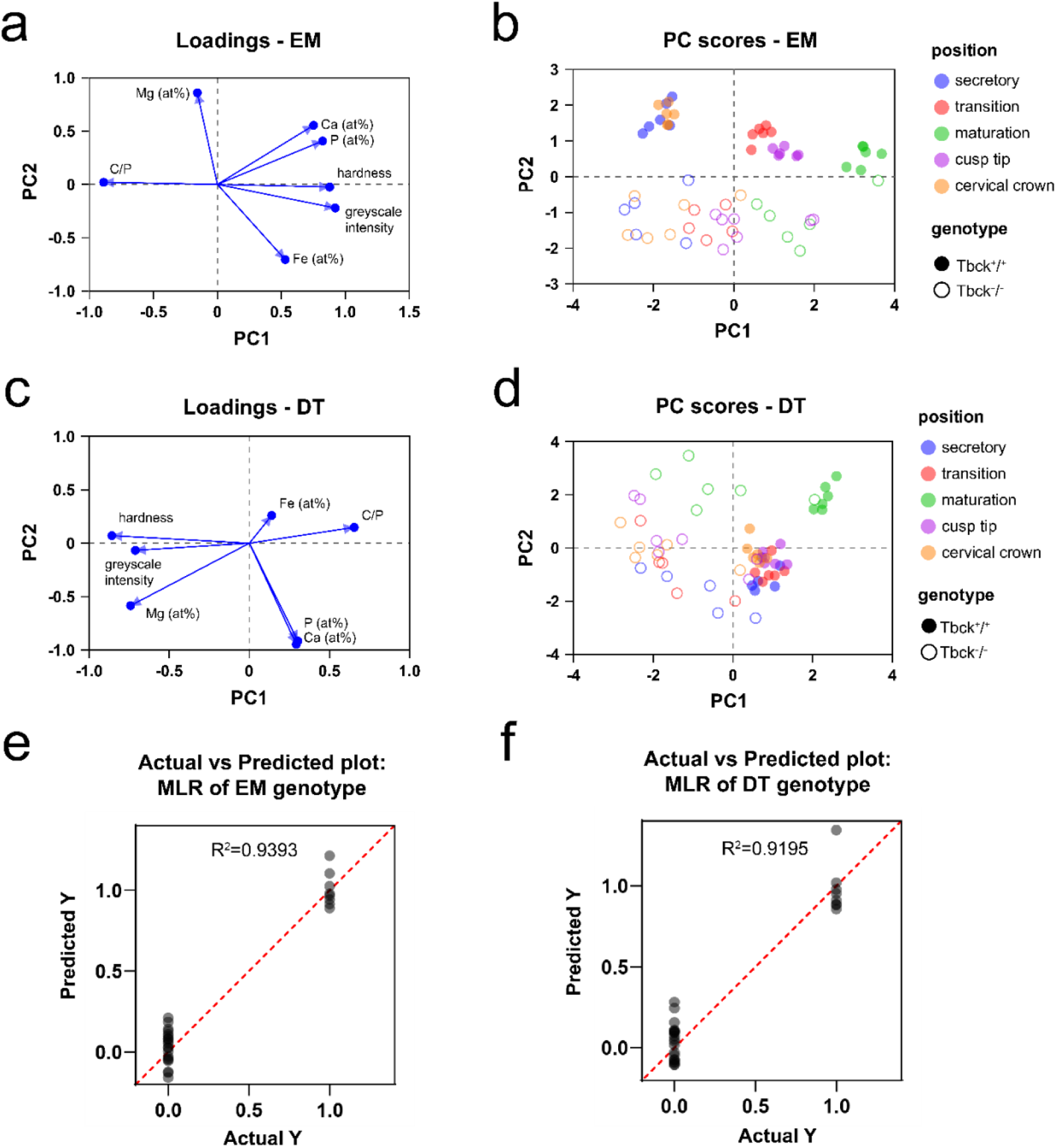
Multimodal integration of microCT, nanoindentation, and compositional metrics by PCA and genotype-predictive MLR. Principal component analysis (PCA) was applied separately to enamel (EM) and dentin (DT) using hardness, EDS-derived Mg, Fe, Ca, and P (at%), microCT greyscale intensity, and the Raman carbonate-to-phosphate ratio (C/P) to visualize coordinated variation across developmental positions and genotype; multiple linear regression (MLR) then leveraged the same inputs to predict Tbck genotype. **a** PCA loadings (EM) showing the contribution and direction of each variable to PC1 and PC2 in enamel. **b** PC scores (EM) for enamel measurements, colored by developmental position (secretory, transition, maturation, cervical crown, cusp tip) and coded by genotype (filled, Tbck^+/+^; open, Tbck^−/−^). **c** PCA loadings (DT) depicting the variable contributions to PC1 and PC2 in dentin. **d** PCA scores (DT) for dentin measurements, colored by position and coded by genotype. **e** MLR genotype prediction (EM) shown as predicted genotype (Y-axis) versus actual genotype (X-axis) for enamel, with the red dashed line indicating perfect agreement (y = x). **f** MLR genotype prediction (DT) shown as predicted versus actual genotype for dentin, with the red dashed line indicating perfect agreement.

PCA reduced the dataset into orthogonal components that successively maximized variance. In enamel, PC1 explained 56.52% of the variance (Eigenvalue 3.956), and PC2 explained 25.12% (Eigenvalue 1.758), together capturing 81.64% of total variance (Table S3). Enamel loadings indicated positive contributions to PC1 from Ca (at%), P (at%), hardness, and greyscale intensity, whereas C/P loaded negatively; Mg (at%) and Fe (at%) contributed primarily along PC2 (Fig. 5a). In the enamel score plot, samples distributed along PC1 by developmental position, while genotype separation was most apparent along PC2 (Fig. 5b). *Tbck* KO enamel also showed an overall shift toward more negative PC1 values relative to WT.

In dentin, variance was distributed across three components, with PC1 accounting for 34.56% (Eigenvalue 2.419) and PC2 accounting for 31.10% (Eigenvalue 2.177); inclusion of PC3 increased the cumulative variance explained to 80.50% (Eigenvalue 1.039) (SI Appendix, Table S4). For visualization, DT score and loading plots are shown for PC1–PC2 (Fig. 5c–d). Dentin loadings suggested that C/P ratio contributed strongly to separation along PC1, whereas Mg (at%), hardness, and greyscale intensity loaded in the opposite direction; Fe (at%), Ca (at%) and P (at%) contributed more prominently to variation along PC2 (Fig. 5c). In the dentin score plot, genotype separation was partially apparent along PC1, whereas developmental position was distributed along PC2, with samples spanning from negative to positive PC2 values across positions (Fig. 5d).

Sex-based stratification did not alter enamel clustering patterns (Fig. S9a). In dentin, *Tbck* WT samples showed minimal separation by sex, whereas female KO dentin exhibited a modest shift toward more negative PC1 values relative to male KO dentin (Fig. S9b).

Building on the PCA structure, we next used MLR to test whether the same integrated variables predict *Tbck* genotype. Separate models were fit for enamel and dentin (Fig. 5e–f). For enamel, predicted values closely matched observed genotype labels (R² = 0.9393; Fig. 5e), and the model was significant by ANOVA (F(9,34) = 58.42, p < 0.0001; Table S3). EDS/Fe (at%) was the most significant predictor (β = 2.971, p < 0.0001), with additional contributions from Ca (at%), Mg (at%), and the C/P ratio. For dentin, the genotype model similarly showed strong agreement between predicted and observed values (R² = 0.9195; Fig. 5f) and was significant by ANOVA (F(9,34) = 43.13, p < 0.0001; Table S4). In dentin, genotype prediction was primarily driven by EDS/Fe (at%) (β = 3.443, p < 0.0001) and EDS/Ca (at%) (β = −0.1336, p = 0.0006) (Table S4).

## Discussion

In this study, we identify a previously unrecognized craniofacial hard-tissue phenotype associated with TBCK deficiency, characterized by alterations in enamel and bone mineralization at mechanical, elemental, and molecular levels. Previous clinical and experimental studies of TBCK syndrome have primarily focused on neurodevelopmental impairment and skeletal fragility, with limited investigation into the quality of mineralized tissues themselves^3–12,16^. By applying a multimodal analytical framework to a *Tbck* knockout mouse model, our findings particularly highlight the enamel signature in genetic development as a sensitive readout of early mineralization perturbations.

A key observation is that TBCK deficiency alters the timing and spatial progression of mineralization rather than producing uniform reductions in mineral density. MicroCT analyses revealed subtle changes in incisor length and eruption-associated grayscale profiles, a common phenotype in monogenic studies ^21–24^, with divergence emerging at the onset of eruption.

Meanwhile, molar mineral density and volume were largely preserved. This pattern is consistent with prior studies demonstrating that developmental disturbances in mineralized tissues often manifest as delays or desynchronization of mineral accrual rather than absolute hypomineralization, particularly in continuously growing structures such as the mouse incisor ^25–27^. The preservation of bulk grayscale intensity reinforces extensive literature showing that microCT is relatively insensitive to changes in mineral quality, crystallinity, or matrix^28,29^.

Histological analyses revealed that enamel matrix disruption in *Tbck* KO incisors emerges during the transition stage and becomes more pronounced during maturation. This timing aligns with a well-established body of enamel biology literature identifying the transition stage as a critical vulnerability window during which ameloblasts briefly differentiate to a matrix-depositing state to a predominantly matrix-degrading and mineral-depositing state ^30–32^. The localization of matrix irregularities near the dentin–enamel junction suggests impaired matrix processing, clearance, and/or defective quality of the proteins being initially deposited, distinctions that parallel phenotypes reported in models of enamel hypomaturation rather than classical defects in enamel secretion ^33,34^.

Consistent with this interpretation, nanoindentation demonstrated that TBCK deficiency selectively compromises late-stage enamel hardening. Enamel modulus and hardness were preserved during secretory and transition stages but were significantly reduced during maturation and at the molar cusp tip. Stage-restricted reductions in enamel mechanical properties have been reported in multiple contexts where mineral transport, matrix degradation, or crystal growth is disrupted without affecting early matrix secretion^17,35–37^. In contrast, the mechanical properties of dentin were preserved across regions, consistent with the collagen-templated and incremental nature of dentin mineralization^38,39^. The accompanying reduction in alveolar bone modulus and hardness is concordant with skeletal fragility and altered bone mineral quality reported in TBCK syndrome and related disorders^10,18,40^.

Elemental analyses further contextualize these mechanical findings within established mineralization paradigms. Reductions in calcium and phosphorus, together with increased carbon content during the transition stage, are consistent with delayed organic matrix removal and incomplete mineral maturation, a pattern reported in both genetic and environmental models of enamel hypomaturation^41–43^. The observation that magnesium is consistently reduced while iron is elevated beginning at the secretory stage is particularly notable. Magnesium is known to regulate apatite crystal size, solubility, and growth kinetics, whereas iron incorporation has been associated with altered crystal chemistry and oxidative microenvironments^44,45^. The early emergence and persistence of these minor-element alterations suggest that TBCK deficiency perturbs mineral incorporation well before overt changes in stiffness are detectable.

Raman spectroscopy provided complementary evidence that TBCK deficiency disrupts late-stage mineral chemistry and crystal organization. The altered maturation trajectory of the ν1 phosphate band, together with an elevated carbonate-to-phosphate ratio at maturation, is consistent with reduced crystal perfection and increased carbonate substitution—features widely associated with mechanically weaker enamel^33,46^. Shifts in ν2 and ν4 phosphate bands further indicate altered phosphate environments within the apatite lattice^47,48^, reinforcing the conclusion that TBCK deficiency primarily affects mineral refinement rather than initial matrix secretion. The absence of genotype-dependent differences in molar enamel carbonate-to- phosphate ratios suggests that continuously erupting enamel may be particularly sensitive to TBCK-dependent processes, consistent with its prolonged and spatially resolved maturation program^49,50^.

Integration of microCT, mechanical, and compositional data using multivariate analyses revealed a robust, genotype-specific mineral signature. As observed in prior multiscale analyses of mineralized tissues, developmental position accounted for a substantial proportion of variance, reflecting the dominant influence of maturation state^17,51,52^. Notably, genotype separation emerged through coordinated shifts in elemental composition and mineral chemistry, with iron and calcium acting as dominant predictors across tissues. These findings underscore the value of integrative analytical approaches for resolving subtle phenotypes that may be obscured when individual modalities are analyzed in isolation^52,53^.

From a clinical perspective, these findings have implications for TBCK syndrome and other rare genetic disorders with delayed or progressive manifestations. Teeth provide a permanent, non-remodeling archive of early development, and enamel in particular is highly sensitive to disruptions in cellular trafficking, matrix processing, and ion transport^54–56^. The identification of enamel-specific mineral defects in a *Tbck* knockout model suggests that dentition may serve as an accessible biomarker of early developmental perturbations, potentially preceding overt clinical symptoms. This is especially relevant given the elevated risk of feeding difficulties, tooth aspiration, and pulmonary complications reported in children with TBCK syndrome^3,4,7^.

More broadly, this study represents the first application of a comprehensive multimodal hard-tissue analysis framework to a genetic syndrome affecting craniofacial development. By situating TBCK deficiency within the broader context of the literature on enamel maturation, mineral chemistry, and skeletal disease, our findings demonstrate that relying solely on conventional imaging may underestimate the disease burden. This work establishes a foundation for extending similar stage-resolved, materials-based analyses to other rare genetic disorders in which craniofacial and dental phenotypes remain underrecognized^57,58^.

## Materials and methods

### Mice

All animal procedures were approved by the Institutional Animal Care and Use Committee (IACUC) of the Children’s Hospital of Philadelphia and were conducted in accordance with institutional guidelines. TBC1 domain containing kinase; targeted mutation 1b, Helmholtz Zentrum Muenchen GmbH), was backcrossed to C57BL/6J for six generations and then backcrossed to C57BL/6N for an additional six generations to generate the mice used in this study. 3 females and 3 males of each genotype (*Tbck^+/+^, Tbck^+/−,^ and Tbck^−/−^)* were littermates and age-matched to P12^59^. Mice of both sexes were included in the analyses. Mice were housed under a 12 h light/dark cycle with ad libitum access to food and water.

For tissue collection, mice were anesthetized with isoflurane and transcardially perfused with 1× phosphate-buffered saline (PBS) followed by 4% paraformaldehyde (PFA) in PBS. After careful soft tissue removal (degloving) and decapitation, whole skulls were post-fixed in 4% PFA for 24 h at 4 °C with gentle agitation. Specimens were subsequently stored in 1× PBS at 4 °C until further processing.

### MicroCT analysis

Acquisition details: A SCANCO MicroCT 45 machine (70kVP, 7.4um, 900ms) was used to scan full skulls in 70% ethanol. (Make sure this is correct, also not sure how you want to call the ones that were partially separated into upper and lower mandible) Following contouring of scans, DICOM files were exported. First molar teeth and the incisors were then segmented using BounTI^60^ with the following parameters: Initial Threshold: 10000, Target Threshold: 6000, Number of Iterations: 100, and Number of Segments: 9. The 9 segments were manually separated from the 10000 threshold segmentations to the 4 first molar teeth, the 4 incisors, and the rest of the bones and teeth as one segment. This was used as the seed to generate the segmentation. Lastly, the bones and other teeth were removed from the final segmentation, and a smoothing of 3 voxels was applied. To visualize the teeth of interest, volume renderings were produced in Avizo (Thermo Fisher Scientific, MA, USA) with greyscale (proxy for density) values coloured from blue to red, ranging from 5000 to 25000 in terms of the greyscale value. The average greyscale values of each tooth were recorded. This analysis was adapted from^17^.

This analysis was augmented by manually locating and placing 8x8x16 voxel cuboids in incisor enamel/dentin at the secretory, transition, and maturation stages, as well as in the first molar regions at the cervical crown and cusp tip. Localized similarly to the nanoindentation. The size of the cuboid was chosen to fully fit inside the enamel, and, following the previous study, the largest dimension was aligned with the gradient to minimize variation due to positional uncertainty. In addition to the local measurements, the grayscale intensity gradient was investigated across the mature enamel. Enamel was separated by a 18000 greyscale threshold. The separated enamel was then positioned in line with the sagittal axis, and the values in each coronal slice were recorded and reported as a function of the slice number (position in the enamel: slice 0 - location where the enamel grayscale value is equal to the dentine value).

### Histological analysis

*Tbck*^+/+^ and *Tbck*^−/−^ C57BL/6N mice (n = 6 per genotype) were collected at postnatal day 12 and perfusion-fixed with 4% paraformaldehyde (PFA) in phosphate-buffered saline (PBS), followed by post-fixation in 4% PFA at 4 °C for 24 h. Samples were decalcified in 8% EDTA (pH 7.4) at 4°C with gentle agitation for 2 weeks. After decalcification, tissues were dehydrated through a graded ethanol series, cleared in xylene, and embedded in paraffin. Serial sagittal sections (5 μm) were cut on a rotary microtome, mounted on charged glass slides, and dried overnight at 37 °C. Hematoxylin and eosin (H&E) staining was performed by the Children’s Hospital of Philadelphia Pathology Core Facility. Stained sections were imaged using a Keyence microscope at magnifications ranging from 10× to 40×.

### Sample preparation

Following microCT, left mandibles were harvested and dehydrated through a graded ethanol series (50%, 70%, 90%, and 100%; 15 min each). Specimens were then infiltrated in a 1:1 ethanol: resin mixture overnight, embedded in 100% LR White resin (Electron Microscopy Sciences, PA), and polymerized at 65 °C for 48h. Resin blocks were sectioned in the sagittal plane using diamond blades (C.L. Sturkey, PA) on a rotome (Leica) to expose the full incisor and first molar. Sections were subsequently polished using a surface grinder (PetroThin, Buehler, IL), rinsed thoroughly with distilled water, and air-dried prior to analysis.

### Nanoindentation

Mechanical properties of rodent teeth and mandibles were measured using an iMicro nanoindenter (Nanomechanics Inc.) at the University of Pennsylvania. Elastic modulus (E) and indentation hardness (H) were determined using a diamond Berkovich tip to a maximum load of 50 mN. Continuous stiffness measurement (CSM) was used to obtain contact stiffness throughout the loading profile^61^. Specimens were mounted on aluminum pucks with Crystalbond 509 and loaded into the instrument sample tray for testing.

Indentations were performed in six anatomical regions: incisor enamel/dentin at the secretory, transition, and maturation stages; first molar regions at the cervical crown and cusp tip; and alveolar bone adjacent to the first molar. For each incisor and molar region, six measurement sites were tested, and four sites were tested in alveolar bone. Indentation sites were spaced 30 μm apart^17^.

### Energy Dispersive X-ray Spectroscopy (EDS) Analysis^17^

Electron microscopy was performed using an FEI Quanta 600 FEG Mark II environmental scanning electron microscope (ESEM) operated in low-vacuum mode (0.38 Torr) at 15 kV with a working distance of 10 mm. Elemental composition of enamel and dentin was quantified by energy-dispersive X-ray spectroscopy (EDS) using an EDAX detector integrated with the Quanta 600 ESEM. Measurements were acquired without sputter coating. The horizontal field width was set to 300 μm for all EDS acquisitions. For incisors and molars, EDS line scans (100 μm length; 2 μm line width; 0.4 μm step size) were collected from enamel to dentin in each region, yielding ∼260 measurement points per scan. For bone, 75 μm line scans were collected, yielding ∼190 measurement points per scan.

### Raman Spectroscopy

Raman spectra were collected using a Raman–near-field scanning optical microscopy (NSOM) system (NT-MDT Spectrum Instrument Ltd., Ireland) equipped with a 660 nm laser. Fluorescence background was removed by subtracting a seventh-order polynomial fit in OMNIC 8 (Thermo Fisher Scientific, MA). Mineral chemistry was quantified using band-area metrics. The carbonate-to-phosphate ratio was calculated by integrating carbonate- and phosphate-associated bands following rubber band correction (15 iterations) in OPUS 7.5 (Bruker Optics, Ettlingen, Germany). Integration windows were defined as ν_1_(PO_4_^3−^) at 920–980 cm^−1^and ν_3_(CO ^2−^) at 1057–1093 cm^−1^. Peak parameters were additionally obtained by fitting the 1030–1100 cm^−1^ region using two Gaussian–Lorentzian functions in OriginPro 2016b (OriginLab, Northampton, MA).

### Principal Component Analysis and Multiple Linear Regression

Principal component analysis (PCA) was used to examine multivariate relationships between Tbck^+/+^ and Tbck^−/−^. PCA was performed in GraphPad Prism using a dataset comprising six animals (three males and three females). Categorical factors included sex, tissue type, and developmental position, and continuous variables included nanoindentation hardness, EDS Mg (at%), EDS Fe (at%), EDS Ca (at%), EDS P (at%), microCT greyscale intensity, and the carbonate-to-phosphate ratio (C/P). Continuous variables were standardized (mean = 0; SD = 1) prior to analysis to enable comparability across measurement scales. The number of principal components retained was determined by parallel analysis (1000 Monte Carlo simulations), retaining components whose eigenvalues exceeded simulated eigenvalues at the 95% confidence level.

Regression modeling was used to assess whether the integrated variables predict genotype. Separate multiple linear regression (MLR) models were fit for Tbck WT and KO using the same continuous variables as PCA, with sex and developmental positions included as covariates. Models were fit by least squares and evaluated using R-squared values. Multicollinearity was assessed using variance inflation factors (VIF).

### Statistical analysis

All data are presented as mean ± standard deviation (SD). Statistical analyses were performed using GraphPad Prism (v10.2.3; GraphPad Software, Boston, MA) and Origin 2021b (OriginLab, Northampton, MA). To evaluate differences in Tbck WT and KO measurements across developmental regions, two-way analysis of variance (ANOVA) with repeated measures was used. One-way ANOVA was applied to compare measurements of alveolar bone between genotypes within each characterization method. Mixed-effects ANOVA with repeated measures was used when positional measurements were treated as a within-subject factor (e.g., Raman spectroscopy and EDS line-scan analyses). Statistical significance was defined as p < 0.05.

## Acknowledgments

We would like to thank the TBCK families for providing the inspiration for this story. We would also like to thank the lab members of the Goldsby Lab, Bhoj Lab, Ahrens-Nicklas Lab, and Vining Lab for their supportive and collegial environments where much of this work took place. Finally, we would like to thank our collaborators at the University of Pennsylvania Singh Center (Drs. Matthew Bruckman, Jaime Ford, Eric Stach), McKay Orthopaedic Research Laboratory PCMD MicroCT Core (Dr. Wen Sang), Dr. David L. Goldsby, and Dr. Ottman A. Tertuliano for their guidance and connections to key resources for this project. This work was partially supported by the Joseph and Josephine Rabinowitz Award for Excellence in Research from Penn Dental Medicine. This work was partly carried out at the Singh Center for Nanotechnology, supported by the NSF National Nanotechnology Coordinated Infrastructure Program under grant NNCI-2025608. Research reported in this publication was supported in part by a NIDCR Supplement to KK from the National Center for Advancing Translational Sciences of the National Institutes of Health under award number KL2TR001879.

---

**Fig. S1.**
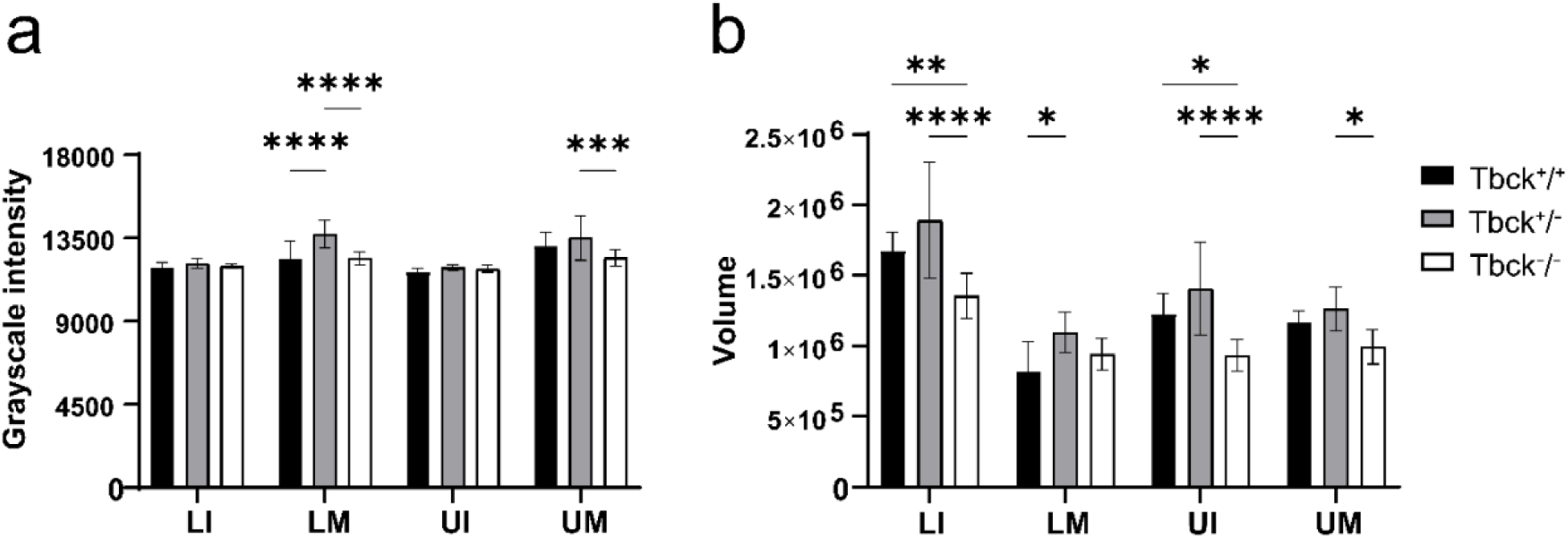
Whole-tooth microCT measurements of mineralization and size across incisors and molars. **a** Mean grayscale intensity of the entire lower incisor (LI), lower molar (LM), upper incisor (UI), and upper molar (UM) for Tbck^+/+^, Tbck^+/−^, and Tbck^−/−^ mice. **b** Total tooth volume for LI, LM, UI, and UM across genotypes. n=6 per genotype. \**p* < 0.05, \*\**p* < 0.01, \*\*\**p* < 0.001, \*\*\*\**p* < 0.0001.

**Fig. S2.**
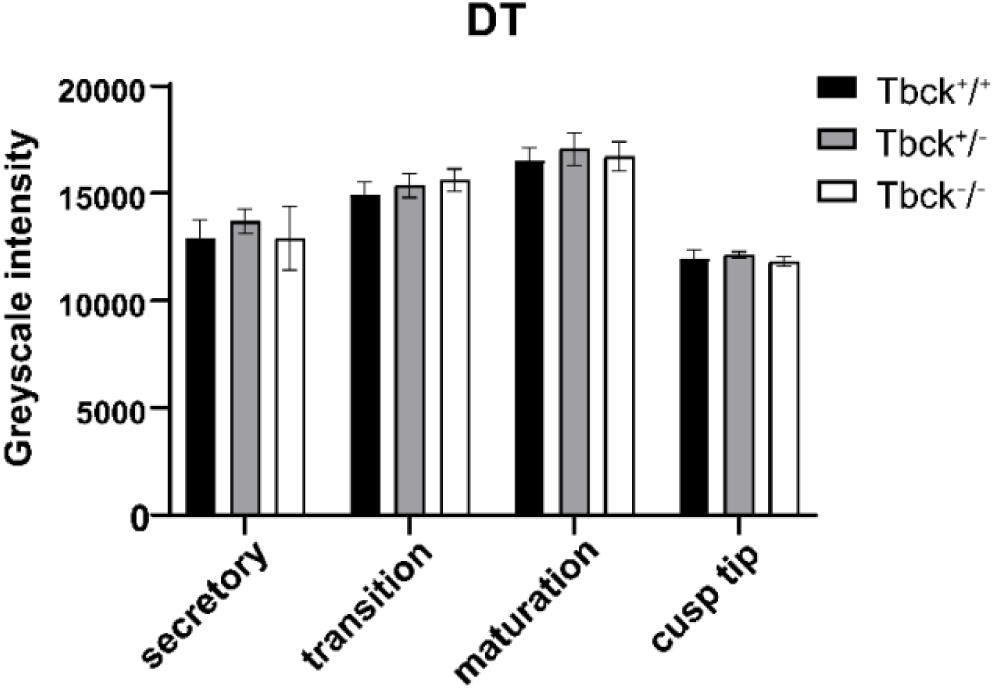
Cropped-region microCT analysis of dentin grayscale intensity. Dentin (DT) grayscale intensity was quantified in cropped regions of interest from incisor (secretory, transition, maturation) and molar dentin (cusp tip) for Tbck^+/+^, Tbck^+/−^, and Tbck^−/−^ mice (n=6).

**Fig. S3.**
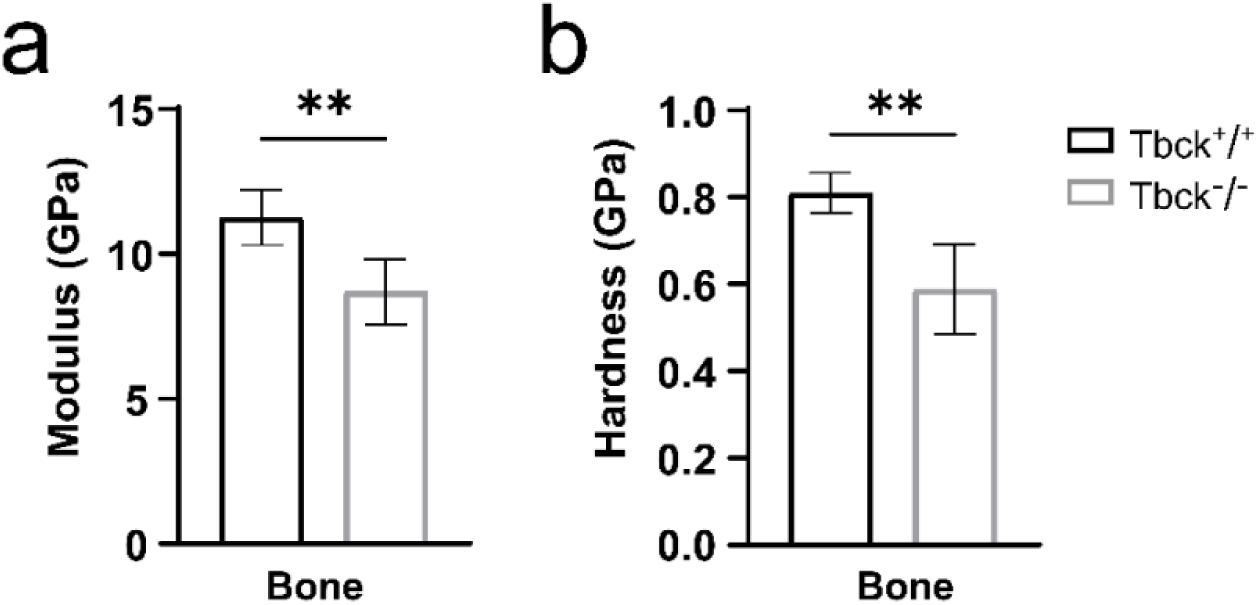
Tbck knockout reduces alveolar bone mechanical properties measured by nanoindentation for **a** modulus and **b** hardness. Data are presented as mean ± SD; mice of both sexes were included (*n* = 4 indent sites per sample). Statistical comparisons were performed using one-way ANOVA; \*\**p* < 0.01.

**Fig. S4.**
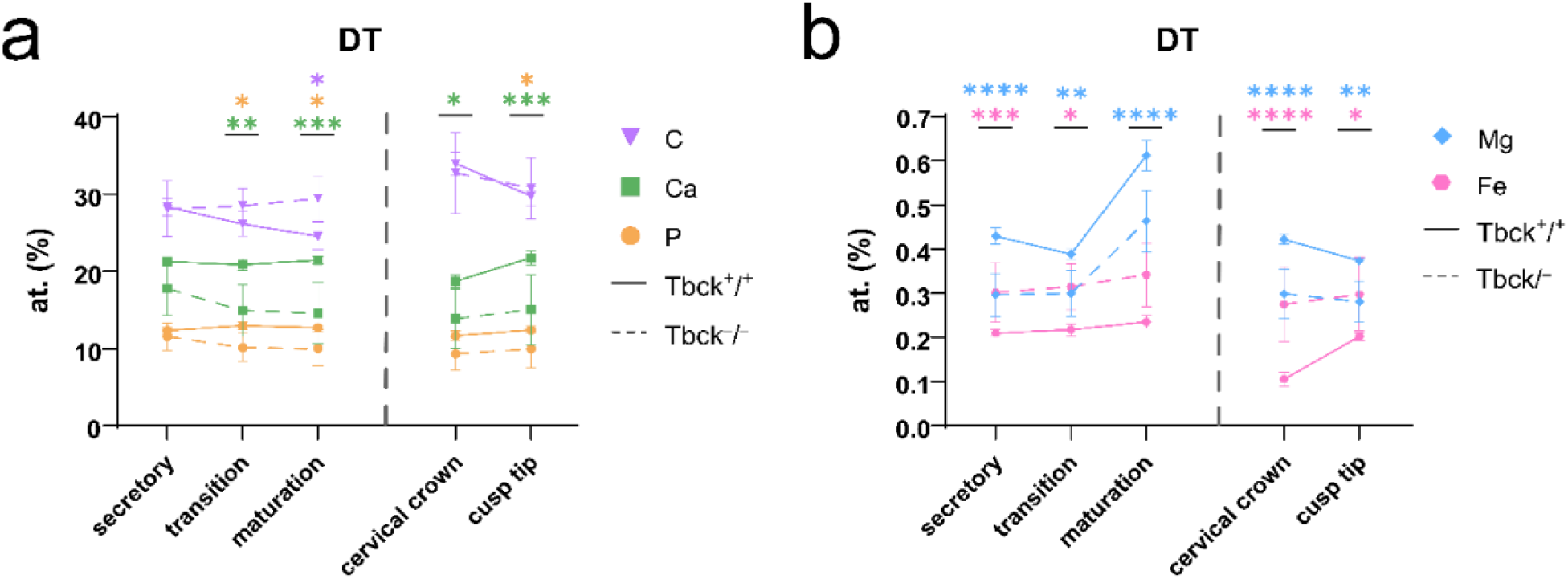
Tbck knockout shifts dentin elemental composition across incisor and molar regions. EDS quantified dentin (DT) elemental profiles in Tbck^+/+^ and Tbck^−/−^ mice, including **a** C, Ca, and P (at%) and **b** Mg and Fe (at%). n=6 per genotype. \**p* < 0.05, \*\**p* < 0.01, \*\*\**p* < 0.001, \*\*\*\**p* < 0.0001.

**Fig. S5.**
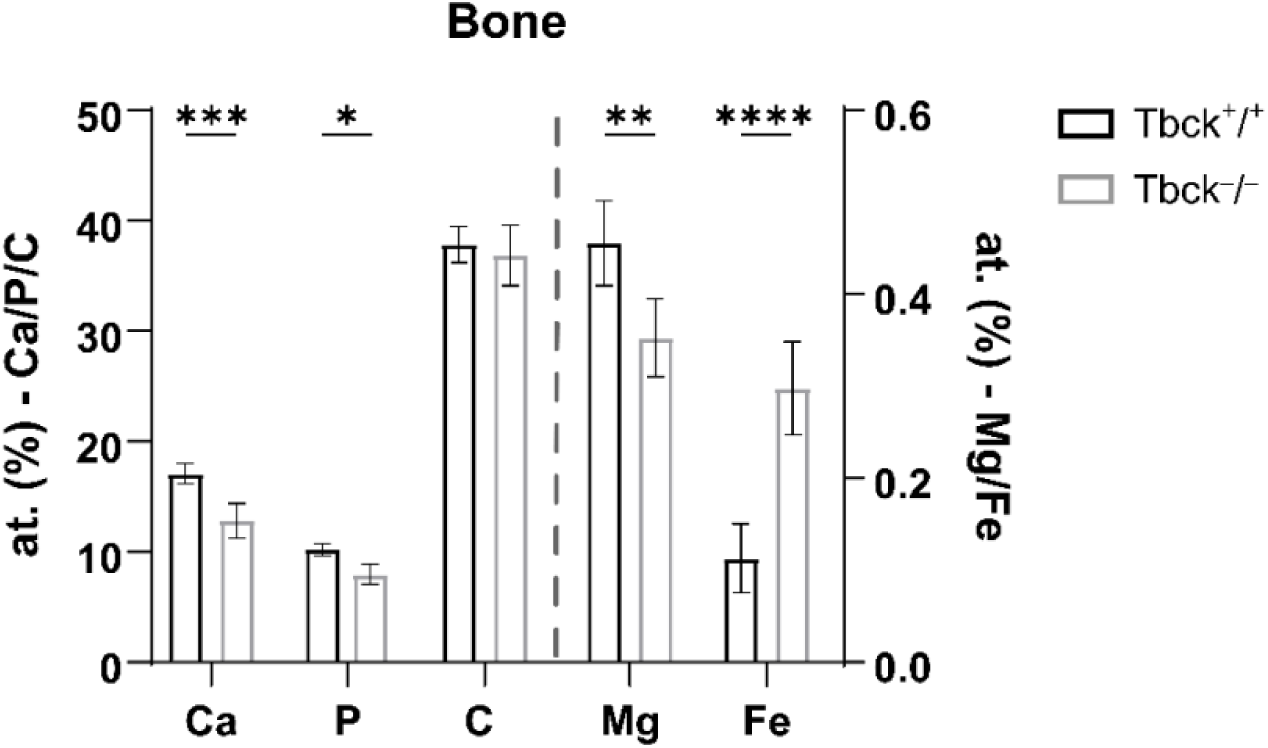
Tbck knockout alters alveolar bone elemental composition. EDS quantified Ca, P, and C (at%, left axis) and Mg and Fe (at%, right axis) in alveolar bone adjacent to the first molar from Tbck^+/+^ and Tbck^−/−^ mice. n=6 per genotype. \**p* < 0.05, \*\**p* < 0.01, \*\*\**p* < 0.001, \*\*\*\**p* < 0.0001.

**Fig. S6.**
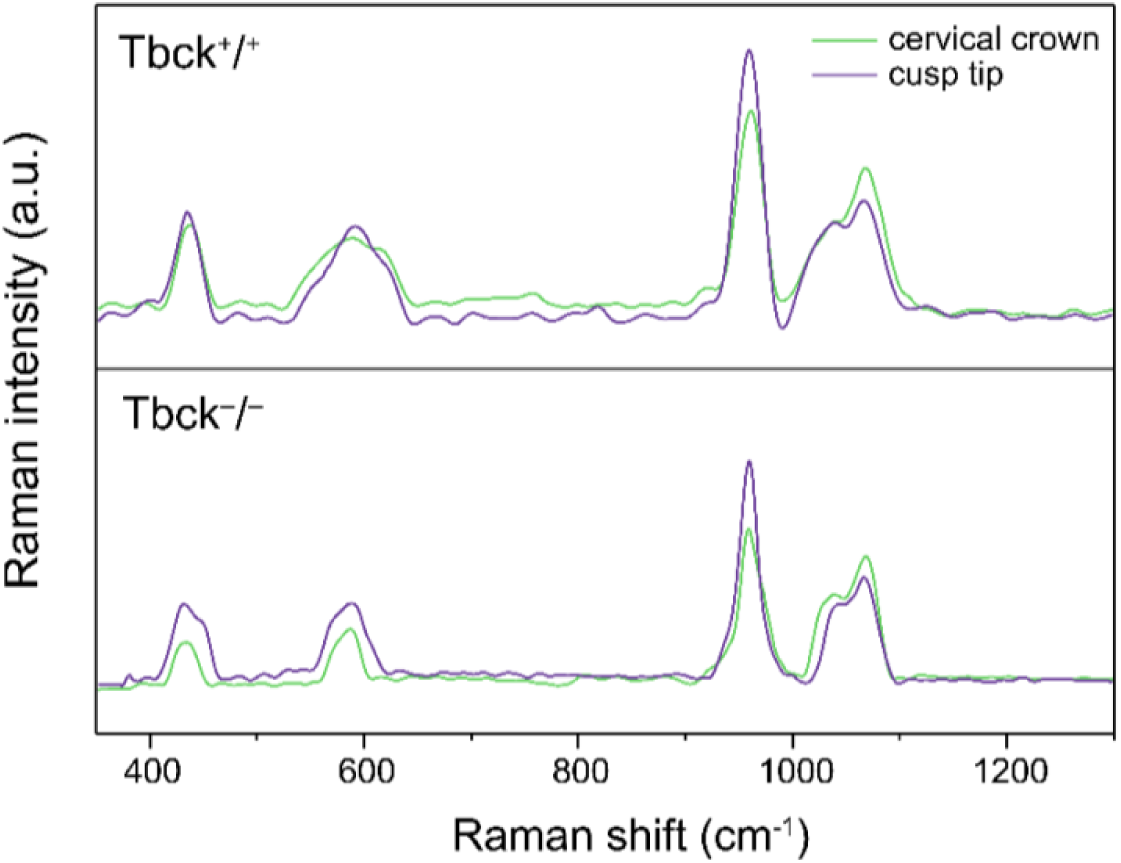
Average Raman spectra from molar enamel at the cervical crown and cusp tip regions in Tbck+/+ (top) and Tbck−/−(bottom) mice (n=6).

**Fig. S7.**
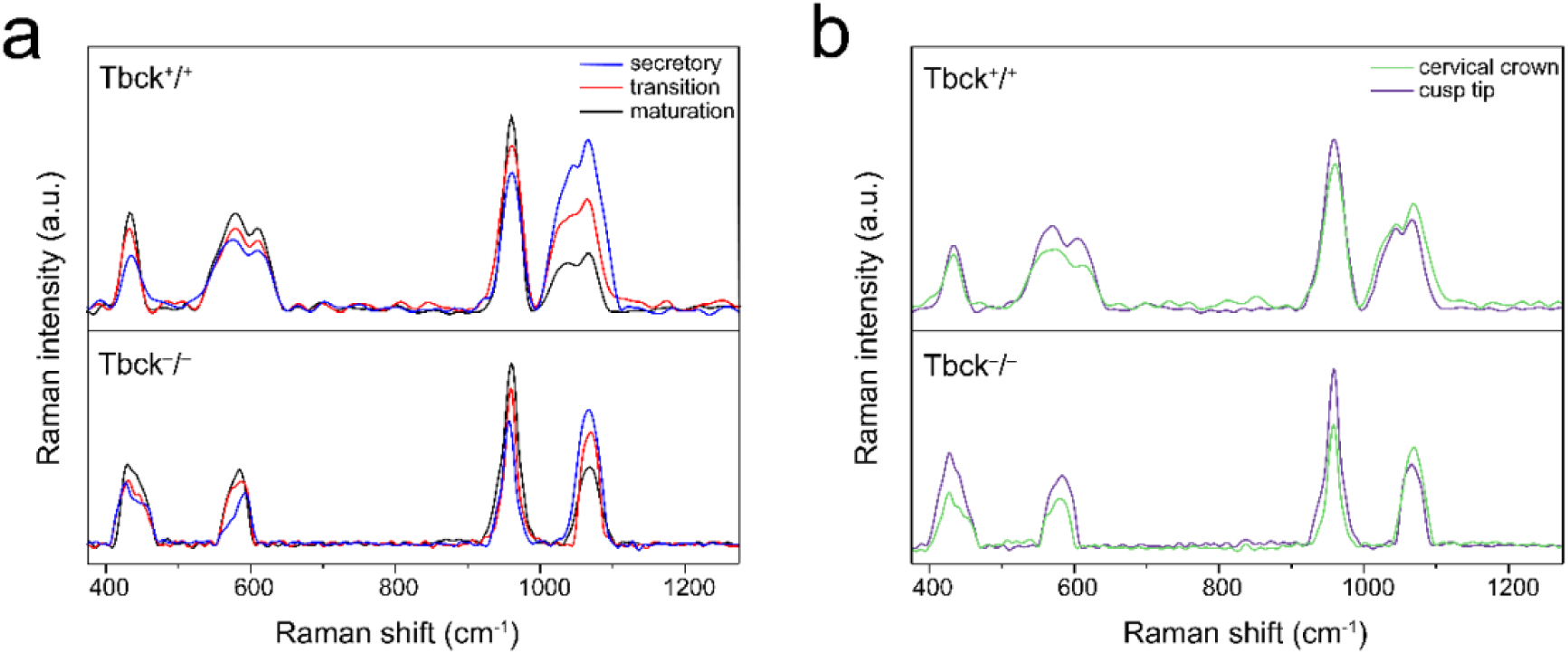
Average Raman spectra from dentin across incisor developmental regions and molar locations in Tbck^+/+^ (top) and Tbck^−/−^(bottom) mice. a Incisor dentin spectra acquired from secretory, transition, and maturation regions. b Molar dentin spectra acquired from the molar cervical crown and cusp tip (n=6).

**Fig. S8.**
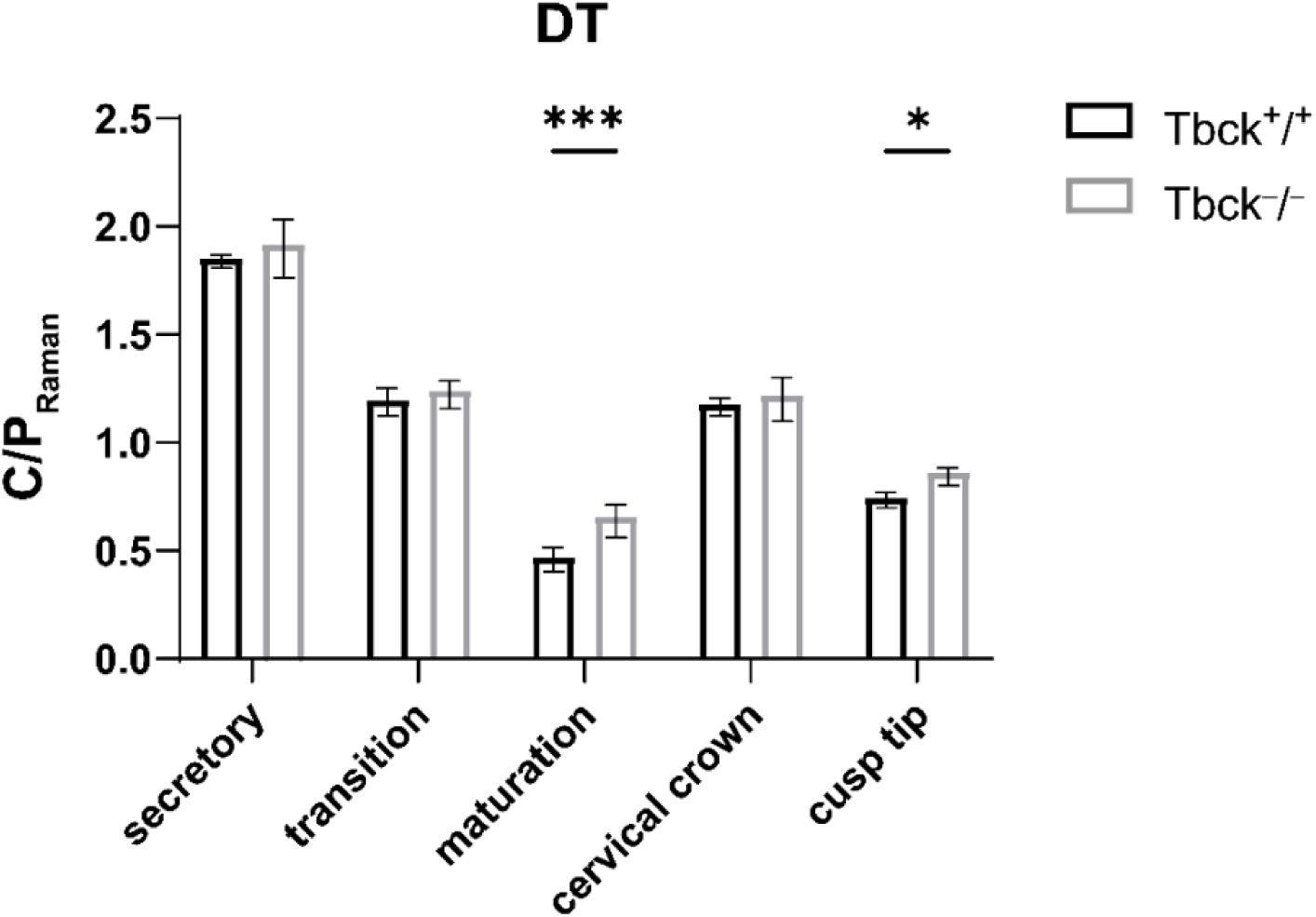
Raman carbonate-to-phosphate ratio in dentin across incisor and molar regions. Statistical comparisons were performed using 2-way ANOVA (n = 6 per genotype); \**p* < 0.05, \*\*\**p*< 0.001.

**Fig. S9.**
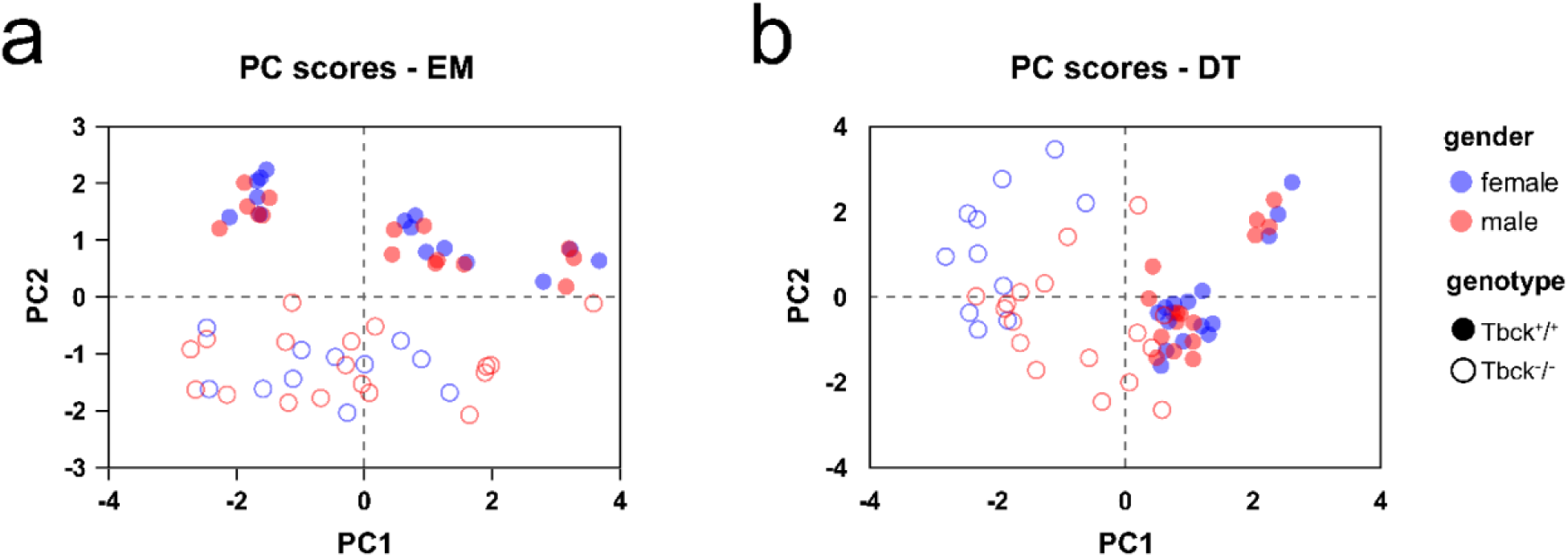
Sex stratification of PCA scores indicates minimal sex-dependent effects on enamel and limited, genotype-restricted effects in dentin. PCA scores are shown for (a) enamel (EM) and (b) dentin (DT), with points colored by sex (female, blue; male, red) and coded by genotype (filled, Tbck^+/+^; open, Tbck^−/−^). **a** EM: Male and female samples overlap within each genotype cluster, indicating no sex-driven separation of enamel scores relative to genotype. **b** DT: WT samples show no sex effect, whereas female KO samples trend toward more negative PC1 compared with male samples.

**Table. S1.** Principal component (PC) summary for enamel (EM) PCA. The table reports eigenvalues and the proportion of variance explained by each component. The first two components were selected for presentation because they capture the majority of variance in the enamel dataset (PC1 = 56.52%, PC2 = 25.12%), together accounting for 81.64% of the total variance.

| Table Analyzed | PCA - EM |  |  |  |  |  |  |
| --- | --- | --- | --- | --- | --- | --- | --- |
| PC summary | PC1 | PC2 | PC3 | PC4 | PC5 | PC6 | PC7 |
| Eigenvalue | 3.956 | 1.758 | 0.6515 | 0.3661 | 0.1445 | 0.1083 | 0.01510 |
| Proportion of variance | 56.52% | 25.12% | 9.31% | 5.23% | 2.06% | 1.55% | 0.22% |
| Cumulative proportion of variance | 56.52% | 81.64% | 90.94% | 96.17% | 98.24% | 99.78% | 100.00% |
| Component selection | **Selected** | **Selected** |  |  |  |  |  |

**Table. S2.** PCA component selection summary for dentin (DT). PC1 and PC2 accounted for 34.56% and 31.10% of variance, respectively (65.66% cumulative), and inclusion of PC3 increased the cumulative variance explained to 80.50%; accordingly, PC1–PC3 were retained for downstream analyses.

| Table Analyzed | PCA - DT |  |  |  |  |  |  |
| --- | --- | --- | --- | --- | --- | --- | --- |
| PC summary | PC1 | PC2 | PC3 | PC4 | PC5 | PC6 | PC7 |
| Eigenvalue | 2.419 | 2.177 | 1.039 | 0.7212 | 0.4900 | 0.1376 | 0.01627 |
| Proportion of variance | 34.56% | 31.10% | 14.84% | 10.30% | 7.00% | 1.97% | 0.23% |
| Cumulative proportion of variance | 34.56% | 65.66% | 80.50% | 90.80% | 97.80% | 99.77% | 100.00% |
| Component selection | **Selected** | **Selected** | **Selected** |  |  |  |  |

**Table. S3.**
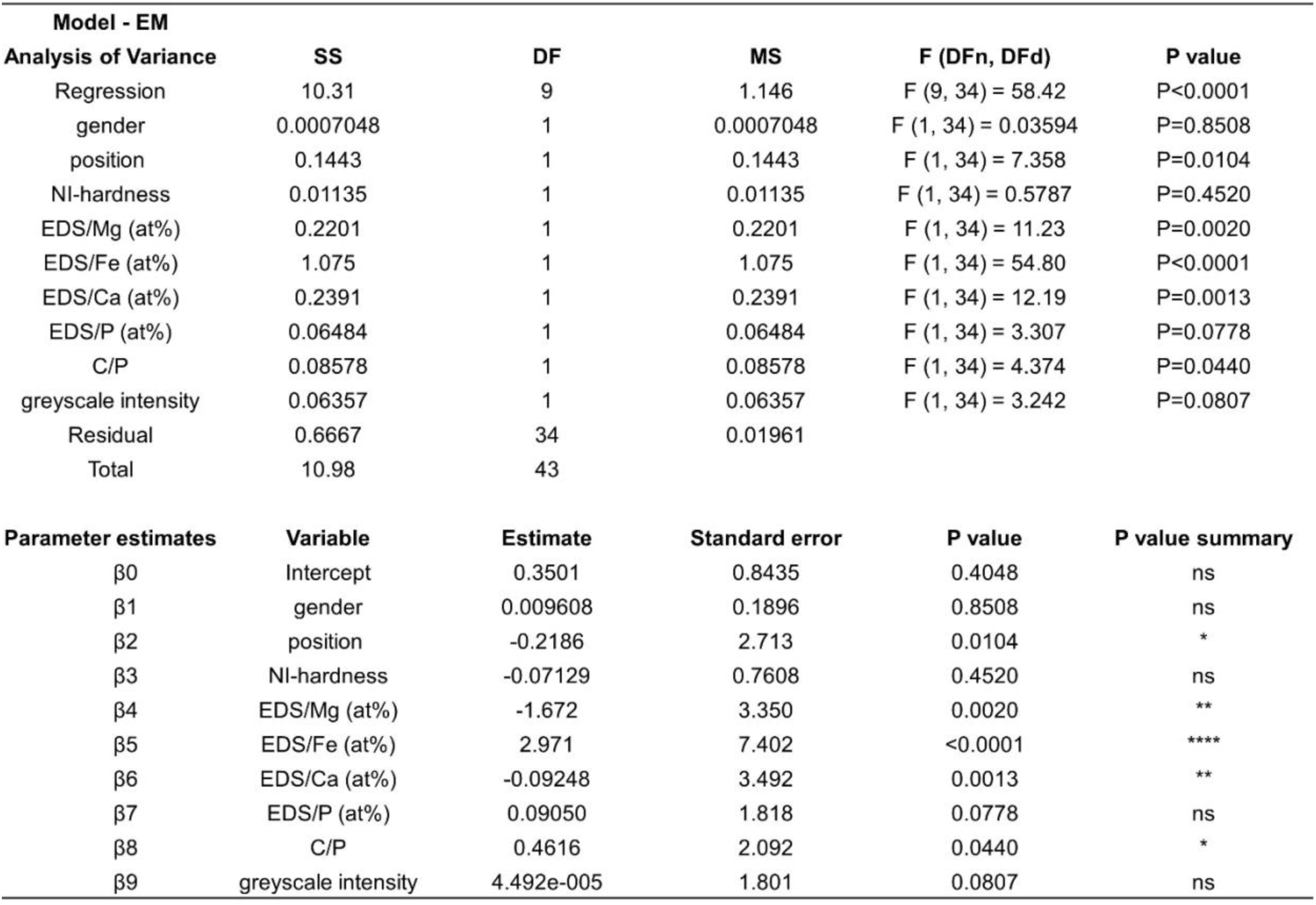
Tabulated outputs from the enamel MLR model, including ANOVA terms (sum of squares, degrees of freedom, mean squares, F-statistics, and p values) and parameter estimates (β), standard errors, and significance for each predictor. The overall model was significant (F(9,34) = 58.42, p < 0.0001). Among predictors, EDS/Fe (at%), EDS/Mg (at%), and EDS/Ca (at%) were significant contributors, with additional effects of developmental position and the Raman-derived carbonate-to-phosphate ratio (C/P), whereas NI-hardness, EDS/P (at%), greyscale intensity, and sex were not significant in the fitted model.

**Table. S4.** Multiple linear regression model for dentin (DT) genotype prediction. The overall model was significant (F(9,34) = 43.13, p < 0.0001). In contrast to enamel, dentin genotype prediction was driven primarily by EDS/Fe (at%) and EDS/Ca (at%), while the other variables were not significant in the fitted model.

| Model - DT |  |  |  |  |  |
| --- | --- | --- | --- | --- | --- |
| Analysis of Variance | SS | DF | MS | F (DFn, DFd) | P value |
| Regression | 10.09 | 9 | 1.121 | F (9, 34) = 43.13 | P<0.0001 |
| gender | 0.05489 | 1 | 0.05489 | F (1, 34) = 2.111 | P=0.1554 |
| position | 1.404e-005 | 1 | 1.404e-005 | F (1, 34) = 0.0005400 | P=0.9816 |
| NI-hardness | 0.05699 | 1 | 0.05699 | F (1, 34) = 2.192 | P=0.1480 |
| EDS/Mg (at%) | 0.04960 | 1 | 0.04960 | F (1, 34) = 1.908 | P=0.1762 |
| EDS/Fe (at%) | 1.405 | 1 | 1.405 | F (1, 34) = 54.03 | P<0.0001 |
| EDS/Ca (at%) | 0.3707 | 1 | 0.3707 | F (1, 34) = 14.26 | P=0.0006 |
| EDS/P (at%) | 0.09857 | 1 | 0.09857 | F (1, 34) = 3.791 | P=0.0598 |
| C/P | 0.0001179 | 1 | 0.0001179 | F (1, 34) = 0.004534 | P=0.9467 |
| greyscale intensity | 0.01138 | 1 | 0.01138 | F (1, 34) = 0.4377 | P=0.5127 |
| Residual | 0.8840 | 34 | 0.02600 |  |  |
| Total | 10.98 | 43 |  |  |  |

**Table. S4** Multiple linear regression model for dentin (DT) genotype prediction.
| Parameter estimates | Variable | Estimate | Standard error | P value | P value summary |
| --- | --- | --- | --- | --- | --- |
| β0 | Intercept | 0.8037 | 0.7409 | 0.4638 | ns |
| β1 | gender | 0.08735 | 1.453 | 0.1554 | ns |
| β2 | position | 0.002732 | 0.02324 | 0.9816 | ns |
| β3 | NI-hardness | 0.4006 | 1.480 | 0.1480 | ns |
| β4 | EDS/Mg (at%) | -0.7633 | 1.381 | 0.1762 | ns |
| β5 | EDS/Fe (at%) | 3.443 | 7.350 | <0.0001 | **** |
| β6 | EDS/Ca (at%) | -0.1336 | 3.776 | 0.0006 | *** |
| β7 | EDS/P (at%) | 0.1331 | 1.947 | 0.0598 | ns |
| β8 | C/P | -0.01860 | 0.06733 | 0.9467 | ns |
| β9 | greyscale intensity | -2.446e-005 | 0.6616 | 0.5127 | ns |

## Reference

1 Smith, A. L., K. & Tomlin, P. D., D.. A familial dysmorphic condition with hypotonia, seizures and precocious puberty. Clin Dysmorphol. 17, 161–164, doi:10.1097/MCD.0b013e328302f0c4 (2008).

2 Alazami, A. M. et al. Accelerating novel candidate gene discovery in neurogenetic disorders via whole-exome sequencing of prescreened multiplex consanguineous families. Cell Rep 10, 148–161, doi:10.1016/j.celrep.2014.12.015 (2015).

3 Zapata-Aldana, E. et al. Further delineation of TBCK - Infantile hypotonia with psychomotor retardation and characteristic facies type 3. Eur J Med Genet 62, 273–277, doi:10.1016/j.ejmg.2018.08.004 (2019).

4 Ortiz-Gonzalez, X. R. et al. Homozygous boricua TBCK mutation causes neurodegeneration and aberrant autophagy. Ann Neurol 83, 153–165, doi:10.1002/ana.25130 (2018).

5 Beck-Wodl, S. et al. Homozygous TBC1 domain-containing kinase (TBCK) mutation causes a novel lysosomal storage disease - a new type of neuronal ceroid lipofuscinosis (CLN15)? Acta Neuropathol Commun 6, 145, doi:10.1186/s40478-018-0646-6 (2018).

6 Guerreiro, R. J. et al. Mutation of TBCK causes a rare recessive developmental disorder. Neurol Genet 2, e76, doi:10.1212/NXG.0000000000000076 (2016).

7 Bhoj, E. J. et al. Mutations in TBCK, Encoding TBC1-Domain-Containing Kinase, Lead to a Recognizable Syndrome of Intellectual Disability and Hypotonia. Am J Hum Genet 98, 782–788, doi:10.1016/j.ajhg.2016.03.016 (2016).

8 Chong, J. X. et al. Recessive Inactivating Mutations in TBCK, Encoding a Rab GTPase-Activating Protein, Cause Severe Infantile Syndromic Encephalopathy. Am J Hum Genet 98, 772–781, doi:10.1016/j.ajhg.2016.01.016 (2016).

9 Mandel, H., Khayat, M., Chervinsky, E., Elpeleg, O. & Shalev, S. TBCK-related intellectual disability syndrome: Case study of two patients. Am J Med Genet A 173, 491–494, doi:10.1002/ajmg.a.38019 (2017).

10 Durham, E. L., et al. TBCK syndrome: a rare multi-organ neurodegenerative disease. Trends Mol Med 29, 783–785, doi:10.1016/j.molmed.2023.06.009 (2023).

11 Tintos-Hernandez, J. A., Santana, A., Keller, K. N. & Ortiz-Gonzalez, X. R. Lysosomal dysfunction impairs mitochondrial quality control and is associated with neurodegeneration in TBCK encephaloneuronopathy. Brain Commun 3, fcab215, doi:10.1093/braincomms/fcab215 (2021).

12 Wu, J. & Lu, G. Multiple functions of TBCK protein in neurodevelopment disorders and tumors. Oncol Lett 21, 17, doi:10.3892/ol.2020.12278 (2021).

13 Moreira, D. P. et al. Neuroprogenitor Cells From Patients With TBCK Encephalopathy Suggest Deregulation of Early Secretory Vesicle Transport. Front Cell Neurosci 15, 803302, doi:10.3389/fncel.2021.803302 (2021).

14 Schuhmacher, J. S. et al. The Rab5 effector FERRY links early endosomes with mRNA localization. Mol Cell 83, 1839–1855 e1813, doi:10.1016/j.molcel.2023.05.012 (2023).

15 Quentin, D. et al. Structural basis of mRNA binding by the human FERRY Rab5 effector complex. Mol Cell 83, 1856–1871 e1859, doi:10.1016/j.molcel.2023.05.009 (2023).

16 Sumathipala, D. et al. TBCK Encephaloneuropathy With Abnormal Lysosomal Storage: Use of a Structural Variant Bioinformatics Pipeline on Whole-Genome Sequencing Data Unravels a 20-Year-Old Clinical Mystery. Pediatr Neurol 96, 74–75, doi:10.1016/j.pediatrneurol.2019.02.001 (2019).

17 Jiang, Y. et al. Multimodal Characterization of Rodent Dental Development. ACS Appl Mater Interfaces 17, 33745–33755, doi:10.1021/acsami.5c08408 (2025).

18 Ortiz-Gonzalez, X. R., Dubbs, H., Keller, K. & Durham, E. L. in GeneReviews (eds M. P. Adam et al.) (University of Washington, Seattle, 2025).

19 Fraser, S. J., Natarajan, A. K., Clark, A. S. S., Drummond, B. K. & Gordon, K. C. A Raman spectroscopic study of teeth affected with molar–incisor hypomineralisation. Journal of Raman Spectroscopy 46, 202–210, doi:10.1002/jrs.4635 (2015).

20 Swietlicka, I. et al. Surface and Structural Studies of Age-Related Changes in Dental Enamel: An Animal Model. Materials (Basel) 15, doi:10.3390/ma15113993 (2022).

21 Zhang, Y. et al. Fluorosed mouse ameloblasts have increased SATB1 retention and Galphaq activity. PLoS One 9, e103994, doi:10.1371/journal.pone.0103994 (2014).

22 Katsura, K. A. et al. WDR72 models of structure and function: a stage-specific regulator of enamel mineralization. Matrix Biol 38, 48–58, doi:10.1016/j.matbio.2014.06.005 (2014).

23 Bronckers, A. L. et al. Composition of mineralizing incisor enamel in cystic fibrosis transmembrane conductance regulator-deficient mice. Eur J Oral Sci 123, 9–16, doi:10.1111/eos.12163 (2015).

24 Nakayama, Y., Holcroft, J. & Ganss, B. Enamel Hypomineralization and Structural Defects in Amelotin-deficient Mice. J Dent Res 94, 697–705, doi:10.1177/0022034514566214 (2015).

25 Jheon, A. H., Seidel, K., Biehs, B. & Klein, O. D. From molecules to mastication: the development and evolution of teeth. Wiley Interdiscip Rev Dev Biol 2, 165–182, doi:10.1002/wdev.63 (2013).

26 Juuri, E. et al. Sox2+ stem cells contribute to all epithelial lineages of the tooth via Sfrp5+ progenitors. Dev Cell 23, 317–328, doi:10.1016/j.devcel.2012.05.012 (2012).

27 Seidel, K. et al. Resolving stem and progenitor cells in the adult mouse incisor through gene co-expression analysis. Elife 6, doi:10.7554/eLife.24712 (2017).

28 Ghavami-Lahiji, M., Davalloo, R. T., Tajziehchi, G. & Shams, P. Micro-computed tomography in preventive and restorative dental research: A review. Imaging Sci Dent 51, 341–350, doi:10.5624/isd.20210087 (2021).

29 Guntoro, P., Ghorbani, Y., Koch, P.-H. & Rosenkranz, J. X-ray Microcomputed Tomography (µCT) for Mineral Characterization: A Review of Data Analysis Methods. Minerals 9, doi:10.3390/min9030183 (2019).

30 Bartlett, J. D. Dental enamel development: proteinases and their enamel matrix substrates. ISRN Dent 2013, 684607, doi:10.1155/2013/684607 (2013).

31 Hu, J. C. et al. Enamelin is critical for ameloblast integrity and enamel ultrastructure formation. PLoS One 9, e89303, doi:10.1371/journal.pone.0089303 (2014).

32 Simmer, J. P., Richardson, A. S., Hu, Y. Y., Smith, C. E. & Ching-Chun Hu, J. A post-classical theory of enamel biomineralization… and why we need one. Int J Oral Sci 4, 129–134, doi:10.1038/ijos.2012.59 (2012).

33 Hu, J. C., Chun, Y. H., Al Hazzazzi, T. & Simmer, J. P. Enamel formation and amelogenesis imperfecta. Cells Tissues Organs 186, 78–85, doi:10.1159/000102683 (2007).

34 Lacruz, R. S., Habelitz, S., Wright, J. T. & Paine, M. L. Dental Enamel Formation and Implications for Oral Health and Disease. Physiol Rev 97, 939–993, doi:10.1152/physrev.00030.2016 (2017).

35 Baldassarri, M., Margolis, H. C. & Beniash, E. Compositional determinants of mechanical properties of enamel. J Dent Res 87, 645–649, doi:10.1177/154405910808700711 (2008).

36 Pugach, M. K. et al. M180 amelogenin processed by MMP20 is sufficient for decussating murine enamel. J Dent Res 92, 1118–1122, doi:10.1177/0022034513506444 (2013).

37 Lacruz, R. S. et al. Targeted overexpression of amelotin disrupts the microstructure of dental enamel. PLoS One 7, e35200, doi:10.1371/journal.pone.0035200 (2012).

38 Moradian-Oldak, J. & George, A. Biomineralization of Enamel and Dentin Mediated by Matrix Proteins. J Dent Res 100, 1020–1029, doi:10.1177/00220345211018405 (2021).

39 Goldberg, M., Kulkarni, A. B., Young, M. & Boskey, A. Dentin: Structure, Composition and Mineralization: The role of dentin ECM in dentin formation and mineralization. Front Biosci 3, 711–735, doi:10.2741/e281 (2012).

40 Nyman, J. S., Granke, M., Singleton, R. C. & Pharr, G. M. Tissue-Level Mechanical Properties of Bone Contributing to Fracture Risk. Curr Osteoporos Rep 14, 138–150, doi:10.1007/s11914-016-0314-3 (2016).

41 Maruyama, K., Henmi, A., Okata, H. & Sasano, Y. Analysis of calcium, phosphorus, and carbon concentrations during developmental calcification of dentin and enamel in rat incisors using scanning electron microscopy with energy dispersive X-ray spectroscopy (SEM-EDX). J Oral Biosci 58, 173–179, doi:10.1016/j.job.2016.08.003 (2016).

42 Hsu, Y.-H. et al. The Characterization of Newly Secreted Dental Enamel by Electron Energy Loss Spectroscopy. Microscopy and Microanalysis 30, doi:10.1093/mam/ozae044.981 (2024).

43 Smith, C. E. L. et al. A Fourth KLK4 Mutation Is Associated with Enamel Hypomineralisation and Structural Abnormalities. Front Physiol 8, 333, doi:10.3389/fphys.2017.00333 (2017).

44 Guggenbuhl, P., Filmon, R., Mabilleau, G., Basle, M. F. & Chappard, D. Iron inhibits hydroxyapatite crystal growth in vitro. Metabolism 57, 903–910, doi:10.1016/j.metabol.2008.02.004 (2008).

45 Kis, V. K. et al. Magnesium incorporation into primary dental enamel and its effect on mechanical properties. Acta Biomater 120, 104–115, doi:10.1016/j.actbio.2020.08.035 (2021).

46 Sa, Y. et al. Compositional, structural and mechanical comparisons of normal enamel and hypomaturation enamel. Acta Biomater 10, 5169–5177, doi:10.1016/j.actbio.2014.08.023 (2014).

47 de Mul, F. F. M., Otto, C. & Greve, J. Calculation of the Raman Line Broadening on Carbonation in Synthetic Hydroxyapatite *Journal of Raman Spectroscopy* **19**, 13–21, doi:10.1002/jrs.1250190104 (1988).

48 Penel, G., Leroy, G., Rey, C. & Bres, E. MicroRaman Spectral Study of the PO4 and CO3 Vibrational Modes in Synthetic and Biological Apatites. Calcified Tissue International 63, 475–481, doi:10.1007/s002239900561 (1998).

49 Goldberg, M., Kellermann, O., Dimitrova-Nakov, S., Harichane, Y. & Baudry, A. Comparative studies between mice molars and incisors are required to draw an overview of enamel structural complexity. Front Physiol 5, 359, doi:10.3389/fphys.2014.00359 (2014).

50 Bui, A. T. et al. Identification of stages of amelogenesis in the continuously growing mandiblular incisor of C57BL/6J male mice throughout life using molar teeth as landmarks. Front Physiol 14, 1144712, doi:10.3389/fphys.2023.1144712 (2023).

51 Kochetkova, T. et al. Composition and micromechanical properties of the femoral neck compact bone in relation to patient age, sex and hip fracture occurrence. Bone 177, 116920, doi:10.1016/j.bone.2023.116920 (2023).

52 Dauphin, Y. Using Microstructures and Composition to Decipher the Alterations of Rodent Teeth in Modern Regurgitation Pellets—A Good News-Bad News Story. Minerals 10, doi:10.3390/min10010063 (2020).

53 Dejea, H. et al. Multi-scale characterization of the spatio-temporal interplay between elemental composition, mineral deposition and remodelling in bone fracture healing. Acta Biomater 167, 135–146, doi:10.1016/j.actbio.2023.06.031 (2023).

54 Wang, S. et al. STIM1 and SLC24A4 Are Critical for Enamel Maturation. J Dent Res 93, 94S–100S, doi:10.1177/0022034514527971 (2014).

55 Davis, K. A. et al. Teeth as Potential New Tools to Measure Early-Life Adversity and Subsequent Mental Health Risk: An Interdisciplinary Review and Conceptual Model. Biol Psychiatry 87, 502–513, doi:10.1016/j.biopsych.2019.09.030 (2020).

56 Katsura, K. et al. WDR72 regulates vesicle trafficking in ameloblasts. Sci Rep 12, 2820, doi:10.1038/s41598-022-06751-1 (2022).

57 Iwata, E., Sah, S. K., Chen, I. P. & Reichenberger, E. Dental abnormalities in rare genetic bone diseases: Literature review. Clin Anat 37, 304–320, doi:10.1002/ca.24117 (2024).

58 Salerno, C. et al. Rare Genetic Syndromes and Oral Anomalies: A Review of the Literature and Case Series with a New Classification Proposal. Children (Basel) 9, doi:10.3390/children9010012 (2021).

59 Nair, D. et al. Heterozygous variants in TBCK cause a mild neurologic syndrome in humans and mice. Am J Med Genet A 191, 2508–2517, doi:10.1002/ajmg.a.63320 (2023).

60 Didziokas, M., Pauws, E., Kolby, L., Khonsari, R. H. & Moazen, M. BounTI (boundary-preserving threshold iteration): A user-friendly tool for automatic hard tissue segmentation. J Anat 245, 829–841, doi:10.1111/joa.14063 (2024).

61 Oliver, W. C. & Pharr, G. M. An improved technique for determining hardness and elastic modulus using load and displacement sensing indentation experiments. Journal of Materials Research 7, 1564–1583, doi:10.1557/jmr.1992.1564 (1992).

